# Proximity labeling reveals spatial regulation of the anaphase-promoting complex/cyclosome by a microtubule adaptor

**DOI:** 10.1101/2022.01.04.474939

**Authors:** Xiaofu Cao, Adnan Shami Shah, Ethan J. Sanford, Marcus B. Smolka, Jeremy M. Baskin

## Abstract

The anaphase-promoting complex/cyclosome (APC/C) coordinates advancement through mitosis via temporally controlled polyubiquitination of effector proteins. Despite the long-appreciated spatial organization of key events in mitosis mediated largely by cytoskeletal networks, the spatial regulation of APC/C, the major mitotic E3 ligase, is poorly understood. Here, we describe a microtubule-resident protein, PLEKHA5, as an interactor of APC/C and spatial regulator of its activity in mitosis. PLEKHA5 knockdown delayed mitotic progression, causing accumulation of APC/C substrates dependent upon the PLEKHA5–APC/C interaction. To assess the spatial importance of this interaction, we developed microtubule-localized proximity biotinylation tools, which revealed that depletion of PLEKHA5 decreased the extent of APC/C association with the microtubule cytoskeleton. This decreased APC/C microtubule-localization in turn prevented efficient loading of APC/C with its co-activator CDC20, leading to defects in E3 ligase catalytic activity. We propose that PLEKHA5 functions as an adaptor of APC/C that promotes its subcellular localization to microtubules and facilitates its activation by CDC20, thus ensuring the timely turnover of key mitotic APC/C substrates and proper progression through mitosis.

## INTRODUCTION

Progression through mitosis involves spatiotemporal coordination of cytoskeletal and membrane dynamics with changes in protein activities, chromosomal dynamics, and signaling events (Morgan, 2006). A major role of the ubiquitin–proteasome system is to orchestrate cell cycle progression through temporally controlled protein degradation programs (Komander and Rape, 2012; Rape, 2017). The anaphase-promoting complex/cyclosome (APC/C) is the primary E3 ubiquitin ligase controlling mitotic events including G2/M transition, early mitosis, the metaphase–anaphase transition, and mitotic exit (King et al., 1995; Sudakin et al., 1995; Pines, 2011).

APC/C is a 14-subunit complex and conserved member of the Cullin-RING E3 ligase family that mediates K11 and K48-linked polyubiquitination of several proteins involved in mitotic progression (Yau et al., 2017). Its ubiquitin ligase activity requires the association of one of two structurally related co-activators, CDC20 and CDH1, after conformational changes due to phosphorylation on multiple subunits (Kimata et al., 2008; Qiao et al., 2016; Zhang et al., 2016). During early mitosis, CDC20 interacts with APC/C, promoting degradation of prometaphase substrates including the cyclin-dependent kinase inhibitor p21 and Cyclin A (Amador et al., 2007; Geley et al., 2001; Wolthuis et al., 2008). Later, APC/C^CDC20^ mediates ubiquitination of Cyclin B1 and the separase inhibitor securin to relieve spindle checkpoint inhibition and facilitate the metaphase–anaphase transition (King et al., 1995; Zur and Brandeis, 2001). In anaphase, CDH1 starts to replace CDC20 as the co-activator, directing the degradation of later mitotic effector proteins like PLK1 to facilitate mitotic exit, and the DNA replication inhibitor geminin in early G1 (Kramer et al., 2000; Listovsky and Sale, 2013; Lindon and Pines, 2004; McGarry and Kirschner, 1998). The temporal control of APC/C-mediated ubiquitination in these processes is well established, including identities of ubiquitination substrates and accessory factors that modulate E3 ligase activity (Sivakumar and Gorbsky, 2015; Alfieri et al., 2017; Watson et al., 2018). However, there are major open questions concerning the spatial organization of the APC/C (Sivakumar and Gorbsky, 2015). These include an incomplete understanding of its subcellular localizations, what factors influence changes in its localization, and how compartmentalization of APC/C at different subcellular locations affects its activity toward specific substrates.

The microtubule cytoskeleton is a major cellular structure that organizes mitotic events, and indeed, immunostaining has localized APC/C to various locations within the microtubule network, including spindle poles/centrosomes, kinetochores, chromosomes, as well as the prophase nucleus (Tugendreich et al., 1995; Tischer et al., 2021; Acquaviva et al., 2004; Topper et al., 2002; Sivakumar et al., 2014; Kraft et al., 2003; Ban et al., 2007). Further, spatially distinct pools of APC/C substrates (e.g., Cyclin B1, securin) can exhibit different degradation rates whether they are freely diffusing or associated with the microtubule network (Clute and Pines, 1999; Huang and Raff, 1999; Shindo et al., 2012), suggesting differential activities of APC/C at these cellular localizations. Collectively, both APC/C and CDC20, as well as key APC/C^CDC20^ substrates, reside in multiple subcellular locations, including the bulk cytosol, the nucleus during early prophase, and several components of the microtubule cytoskeletal network.

Yet, a full picture of where APC/C is productively loaded with its co-activator CDC20 and catalyzes its ubiquitination reactions, and what factors regulate recruitment to those subcellular locations, is incomplete (Sivakumar and Gorbsky, 2015). This gap in our understanding of the spatial organization of APC/C is partly due to the challenges associated with effectively determining the subcellular localizations of this large, multi-subunit protein complex with conventional imaging methods. Commercially available antibodies for different APC/C subunits have failed to consistently mark the subcellular localizations of the complex by immunofluorescence, and exogenous overexpression of GFP fusions to individual APC/C subunits may not reliably assemble into functional APC/C complexes (Huang and Raff, 2002; Acquaviva et al., 2004). Moreover, a recent high-throughput effort (OpenCell) has generated CRISPR knockin cell lines with GFP fusions to >1300 human proteins expressed endogenously (Cho et al., 2022); yet for several APC/C subunits in this collection, no reliable localization was observed to intracellular organelles or structures.

Fortunately, recent developments of enzyme-mediated proximity labeling techniques such as BioID/TurboID and APEX provide new avenues for mapping the subcellular localizations of untagged, endogenous proteins with near-nanometer spatial resolution (Zhou and Zou, 2020). In these strategies, an organelle-tethered enzyme releases a highly reactive intermediate, typically a biotin derivative, enabling covalent biotinylation of nearby proteins within a limited radius. As such, proximity biotinylation has been widely applied for mapping the proteomes of organelles and other discrete structures such as membrane contact sites and synaptic clefts. Notably, the covalent nature of the labeling allows for sensitive detection and quantification of even minor or transient spatially restricted pools of proteins that exhibit dynamic behavior and/or can exist in multiple subcellular localizations, including large cytosolic pools, all of which are very challenging to visualize with standard imaging-based methods.

Here, using newly developed microtubule-localized proximity biotinylation tools to assess APC/C localization, we reveal a general mechanism by which the spatial organization of APC/C is coordinated to ensure its proper activity during mitosis. First, we identify PLEKHA5 (pleckstrin homology domain-containing family A, member 5) as a microtubule-localized adaptor protein that interacts with APC/C using mass spectrometry-enabled quantitative proteomics. Second, we find that depletion of PLEKHA5 causes (i) delays in mitotic progression, (ii) reductions in the association of APC/C with CDC20 and the E3 ligase activity of APC/C, and (iii) buildup of several APC/C^CDC20^ substrates in a manner dependent upon the PLEKHA5–APC/C interaction. Third, using two microtubule-targeting TurboID constructs, we demonstrate that a pool of APC/C localizes to the microtubule network and that PLEKHA5 knockdown reduces the levels of this microtubule-localized APC/C pool. We propose that PLEKHA5 functions as a microtubule-specific APC/C adaptor to recruit a pool of APC/C to the microtubule cytoskeleton, promoting its loading with CDC20 and enabling the APC/C^CDC20^-mediated ubiquitination of key mitotic effector proteins, thus facilitating mitosis progression. In this model, PLEKHA5 regulates APC/C subcellular localization and thus accelerates its access to a critical co-activator required for its functions in mitosis, effectively reducing a 3D search in the cytosol to a 1D search along microtubule tracks. This model establishes a framework for understanding how spatial organization of the APC/C impacts its ubiquitination activity and roles in promoting mitotic progression. Further, our study establishes a set of “in vivo biochemistry”-based TurboID tools for interrogating the microtubule-associated proteome, highlighting the versatility of proximity labeling methods for examining unresolved questions in cell biology.

## RESULTS

### PLEKHA5 localizes to microtubules via its N-terminal tandem WW domain

We began this study by focusing on the molecular and cellular properties of PLEKHA5, as part of a larger undertaking to explore roles of how putative phosphoinositide-binding proteins might link lipid binding to the regulation of cell signaling, and in particular, ubiquitination pathways. Toward this effort, we have centered on poorly characterized members of the PLEKHA subfamily of pleckstrin homology domain-containing proteins. Previously, we found that PLEKHA4/kramer regulates Wnt signaling pathways in mammalian cells and in *Drosophila* and promotes proliferation in melanoma by antagonizing the polyubiquitination of Dishevelled by the Cullin 3–KLHL12 E3 ligase (Shami Shah et al., 2019, 2021). The PLEKHA4 paralog PLEKHA5 shares a similar multidomain architecture (**Figure 1A**) and has been linked to proliferation and metastasis in several disease models (Jilaveanu et al., 2015; Liu et al., 2020; Nagamura et al., 2021), though underlying mechanisms remained unknown, prompting us to investigate its molecular properties and cellular functions.

**Figure 1.**
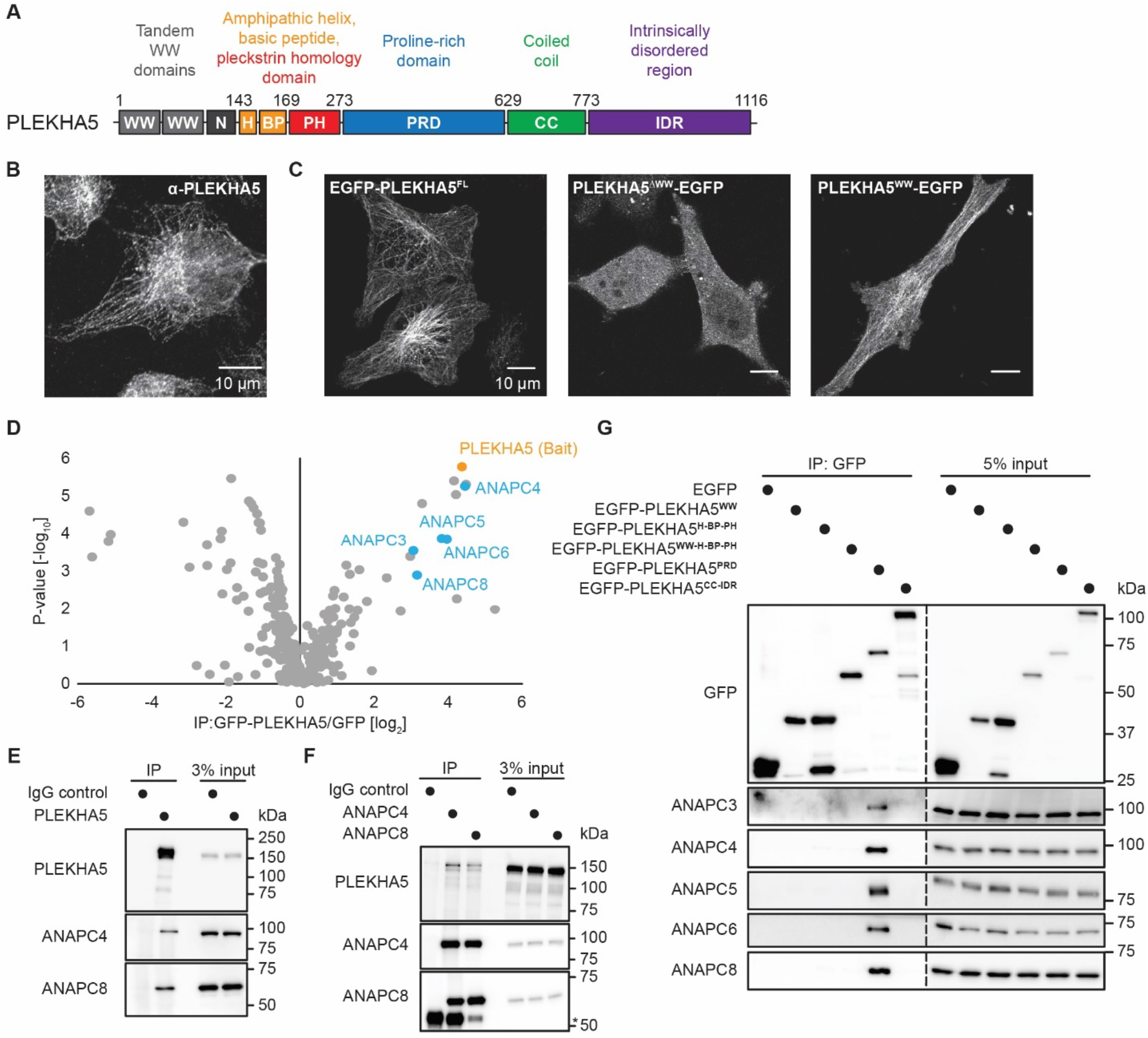
PLEKHA5 localizes to the microtubule network and interacts with several APC/C subunits. (A) Domain map of PLEKHA5. (B) Immunofluorescence (IF) of endogenous PLEKHA5 in HeLa cells by confocal microscopy. (C) PLEKHA5 localizes to the microtubule network via its tandem WW domain. Shown are confocal microscopy of HeLa cells transfected with GFP-tagged full-length protein (EGFP-PLEKHA5^FL^), a truncation lacking the tandem WW domains (PLEKHA5^ΔWW^-EGFP), or the isolated tandem WW domains (PLEKHA5^WW^-EGFP). (D–G) PLEKHA5 interacts with APC/C. (D) A volcano plot from SILAC MS proteomics experiments to determine the PLEKHA5 interactome, where proteins that are statistically significantly enriched in α-GFP immunoprecipitates (IP) from GFP-PLEKHA5-expressing HeLa cells compared to the IP from GFP-expressing cells, including several APC/C subunits, appear on the top right. Statistical significance was determined by a Mann-Whitney U test. (E–F) Western blot analysis of co-IP experiments for endogenous PLEKHA5 (E) and the APC/C subunits ANAPC4 and ANAPC8 (F) to demonstrate the interaction between endogenous PLEKHA5 and the APC/C. IgG heavy chain from the IP is marked by an asterisk. (G) Truncation studies to reveal the PLEKHA5 proline-rich domain (PRD) as sufficient to interact with APC/C. Shown is Western blot analysis of α-GFP IP from HeLa cells transfected with GFP or the indicated GFP-PLEKHA5 fragment. For IF analysis of PLEKHA5 in mitosis, colocalization of GFP-PLEKHA5 with a microtubule marker, and interaction mapping with additional constructs, see Figure S1.

We first examined the cellular localization of PLEKHA5. Unlike PLEKHA4, which localizes to the plasma membrane (Shami Shah et al., 2019), endogenous PLEKHA5 was detected predominantly along the microtubule network in HeLa cells by immunofluorescence (**Figures 1B and S1A**) and imaging of GFP-tagged full-length PLEKHA5 (EGFP-PLEKHA5^FL^, **Figures 1C and S1B**). As a member of the WW-PLEKHA subfamily of proteins (Sluysmans et al., 2021b), PLEKHA5 contains two N-terminal WW domains absent from PLEKHA4. We hypothesized that the microtubule localization of PLEKHA5 might be mediated by its tandem WW domains. Indeed, we found that a GFP-tagged PLEKHA5 construct lacking the two WW domains failed to localize to microtubules, and a GFP fusion to the tandem WW domains localized to microtubules similar to the full-length protein (**Figure 1C**). These data indicate that the tandem WW domain is necessary and sufficient to recruit PLEKHA5 to the microtubule network. These observations are consistent with recent studies demonstrating that the tandem WW domain of PLEKHA5 associates with PDZD11 and that this interaction was crucial for PLEKHA5 to localize to the microtubule network (Sluysmans et al., 2021b; a).

### PLEKHA5 interacts with APC/C, a key E3 ubiquitin ligase regulating mitosis

To explore the molecular functions of PLEKHA5, we identified potential protein-protein interaction partners using stable isotope labeling by amino acids in cell culture (SILAC)-based quantitative mass spectrometry. We performed anti-GFP immunoprecipitation (IP) from HeLa cells stably expressing EGFP-PLEKHA5 or EGFP and subjected the immunoprecipitates to bottom-up proteomics analysis. We identified several known interactors, in agreement with existing datasets, including KCTD3, KRAS, FLOT1, and PDZD11 (**Table S1**; Rolland et al., 2014; Boldt et al., 2016; Go et al., 2021). Interestingly, among the most strongly enriched proteins were several subunits of the anaphase-promoting complex/cyclosome (APC/C) (**Figure 1D, Table S1**).

We validated the interaction of endogenous PLEKHA5 with two representative APC/C subunits, ANAPC4 and ANAPC8 by co-IP followed by Western blotting (**Figures 1E–F**). To map the interacting regions, we performed co-IP of APC/C subunits with several GFP-tagged constructs corresponding to truncated forms of PLEKHA5 or isolated PLEKHA5 domains (**Figures 1G S1C**). The GFP fusion to the proline-rich domain (PRD) (EGFP-PLEKHA5^PRD^) exhibited a strong interaction with APC/C (**Figure 1G**), whereas a construct lacking the PRD (EGFP-PLEKHA5^ΔPRD^), despite its high expression levels, failed to interact with APC/C (**Figure S1C**). These results suggest that the PRD of PLEKHA5 is sufficient and necessary for the interaction with APC/C.

### PLEKHA5 knockdown causes increases in the fraction of cells in G2 and M phases and the levels of mitotic APC/C substrates

The association of PLEKHA5 with APC/C suggests potential functions of PLEKHA5 in mitosis, as APC/C is the primary E3 ligase controlling mitotic entry, progression, and exit. We therefore explored whether the PLEKHA5–APC/C interaction would affect the timing and progression of mitotic events. To avoid deleterious effects on M phase progression and to assess effects in mitotic entry, we first used S phase synchronization and release via double thymidine block (DTB) (Chen and Deng, 2018; Ma and Poon, 2017) and analyzed the cell-cycle phase of HeLa cells transfected with either control or two different PLEKHA5 siRNA duplexes by using propidium iodide staining of fixed cells followed by flow cytometry (**Figure 2A**). We found that depletion of PLEKHA5 by both siRNAs led to an increased percentage of cells retained in G2/M phase after release from DTB (**Figures 2B–D and S2A–C**). We also analyzed the levels of several G2 and M phase protein markers. In HeLa cells synchronized to and then released from S phase, PLEKHA5 knockdown caused a delay in the appearance and the peak expression of a mitosis marker, phospho-histone H3 at serine 10 (H3 S10p) (**Figures 2E–F and S2D–E**). Further, the analysis of CDK1, a major kinase that positively regulates mitosis onset and advance, revealed prolonged expression of the inactive form, which was bearing inhibitory phosphorylation at tyrosine 15 (CDK1 Y15p) (Jin et al., 1996), at later timepoints in PLEKHA5-depleted cells compared to control (**Figures 2E–F and S2D–E**). Additionally, the G2 and M-phase related cyclins (A and B1) showed aberrant accumulation upon PLEKHA5 knockdown (**Figures 2E–F and S2D–E**). Cyclins A and B1 are APC/C substrates in early mitosis and the metaphase-anaphase transition, respectively (Geley et al., 2001; King et al., 1995), so we reasoned that their accumulation in siPLEKHA5 cells might result from a defect in APC/C-mediated polyubiquitination and subsequent proteasomal degradation. Therefore, we examined the expression patterns of two additional APC/C substrates, p21 and securin (Amador et al., 2007; Zur and Brandeis, 2001), in cells released from DTB synchronization. Consistent with our observations for cyclins A and B1, levels of both p21 and securin increased upon PLEKHA5 knockdown (**Figures 2E–F and S2D–E**). These assays demonstrate that depletion of PLEKHA5 antagonizes mitotic entry and progression and suggest that PLEKHA5 may play a role in regulating mitotic events through modulating APC/C.

**Figure 2.**
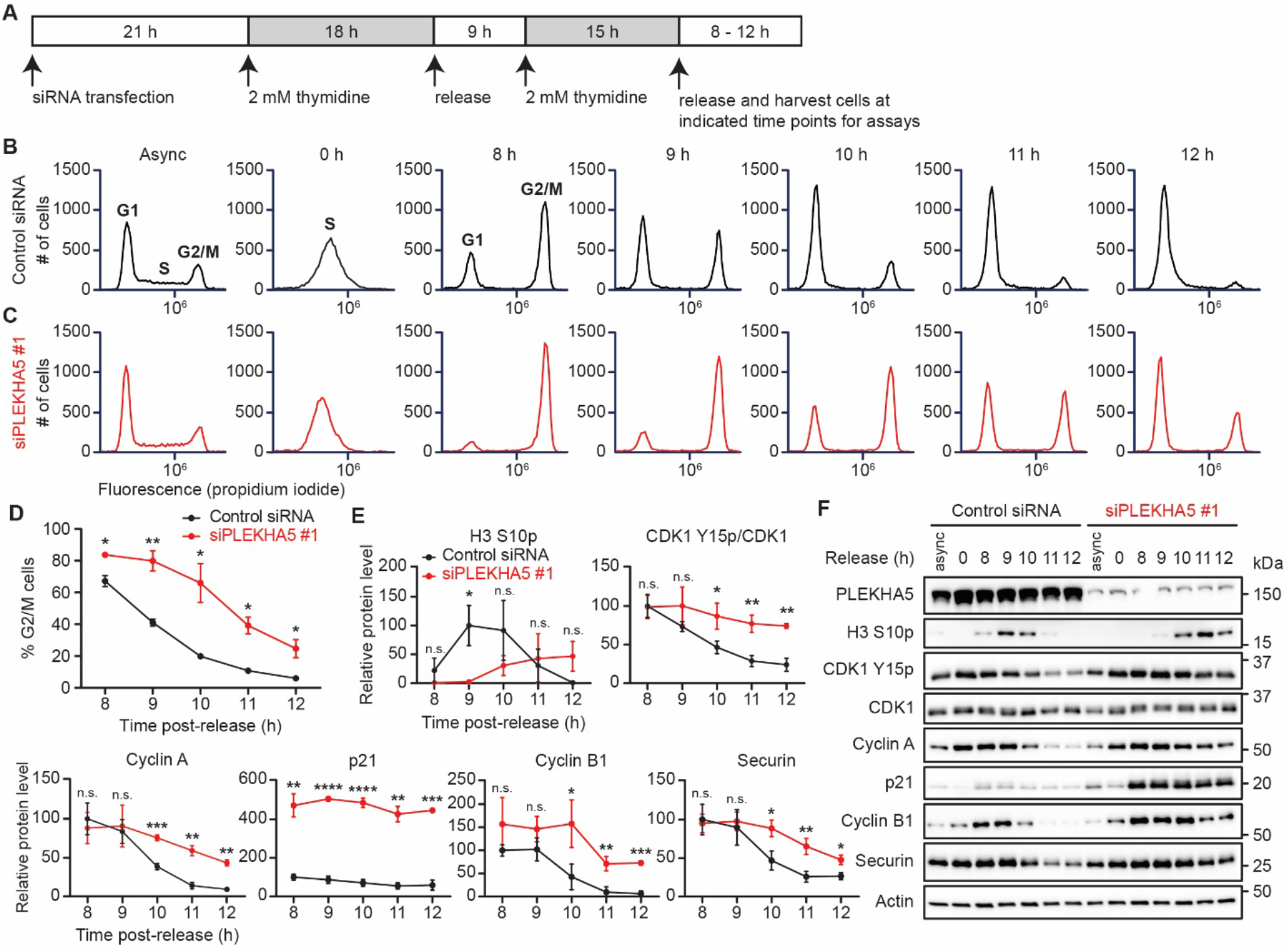
PLEKHA5 knockdown causes an accumulation of cells in G2/M phase and a buildup of APC/C substrates. (A) Schematic representation of experiment timeline. HeLa cells were treated with an siRNA duplex against PLEKHA5 (siPLEKHA5 #1) or a negative control siRNA and synchronized to S phase by a double thymidine block (DTB). Cells were then rinsed, released into fresh growth medium, and harvested at indicated time points after the release. (B–D) siPLEKHA5 #1 treatment leads to accumulation of HeLa cells in G2/M phase. DNA content of asynchronous (async) or DTB-synchronized/released HeLa cells transfected with control siRNA (B) or siPLEKHA5 #1 (C) were evaluated by propidium iodide staining and flow cytometry analysis, and the percentage of cells in G2/M phase were quantified and shown in (D) (n=3). (E– F) G2/M protein markers and APC/C substrates persist in cells treated with siPLEKHA5 #1. Shown are quantification of protein levels (E) and representative Western blots (F) of lysates from asynchronous or DTB-synchronized/released HeLa cells subject to control siRNA or siPLEKHA5 #1 (n=3). Student’s t-test: n.s. not significant; * p < 0.05; ** p < 0.01; *** p<0.001; **** p < 0.0001. For analysis of effects using siPLEKHA5 #2, see Figure S2.

### Interaction of PLEKHA5 with APC/C is required to rescue mitotic defects caused by PLEKHA5 knockdown

To further investigate the mechanism of the PLEKHA5–APC/C interaction in regulating mitosis, we performed rescue experiments with siPLEKHA5 #1–resistant EGFP, EGFP-PLEKHA5^FL^, and EGFP-PLEKHA5^δPRD^ constructs (**Figure 3A**). We confirmed that EGFP-PLEKHA5^ΔPRD^ still localized correctly to the microtubule network (**Figure 3B**). In accordance with previous results, control rescue (with EGFP) in siPLEKHA5-treated cells led to a significant increase of G2 and M-phase protein markers and APC/C substrates (**Figures 3C–D**, compare the first and second bands in groups of four in representative blots, e.g., lanes 9 vs. 10, and the black and red line in quantification plots, significance marked by *). Importantly, the changes in protein expression patterns induced by PLEKHA5 depletion were partially restored by introduction of an siRNA-resistant form of full-length PLEKHA5 protein (**Figures 3C–D**, compare the second and third bands in groups of four in representative blots, e.g., lanes 10 vs. 11, and the red and dark green line in quantification plots, significance marked by #). For example, the peak expression of H3 S10p shifted earlier, to 9 h post-release, and its levels decreased to a greater extent at later time points. Similarly, the buildup of other markers and APC/C substrates were partially reduced by expression of EGFP-PLEKHA5^FL^ in cells treated with siPLEKHA5. However, rescue was not observed in cells expressing siRNA-resistant EGFP-PLEKHA5^δPRD^, a construct unable to interact with APC/C (**Figures S1C and 3C–D**, compare the second and fourth bands in groups of four in representative blots, e.g., lanes 10 vs. 12, and the red and light green line in quantification plots, significance marked by †). These rescue experiments reveal that the PLEKHA5–APC/C interaction is critical for proper advancement of mitosis.

**Figure 3.**
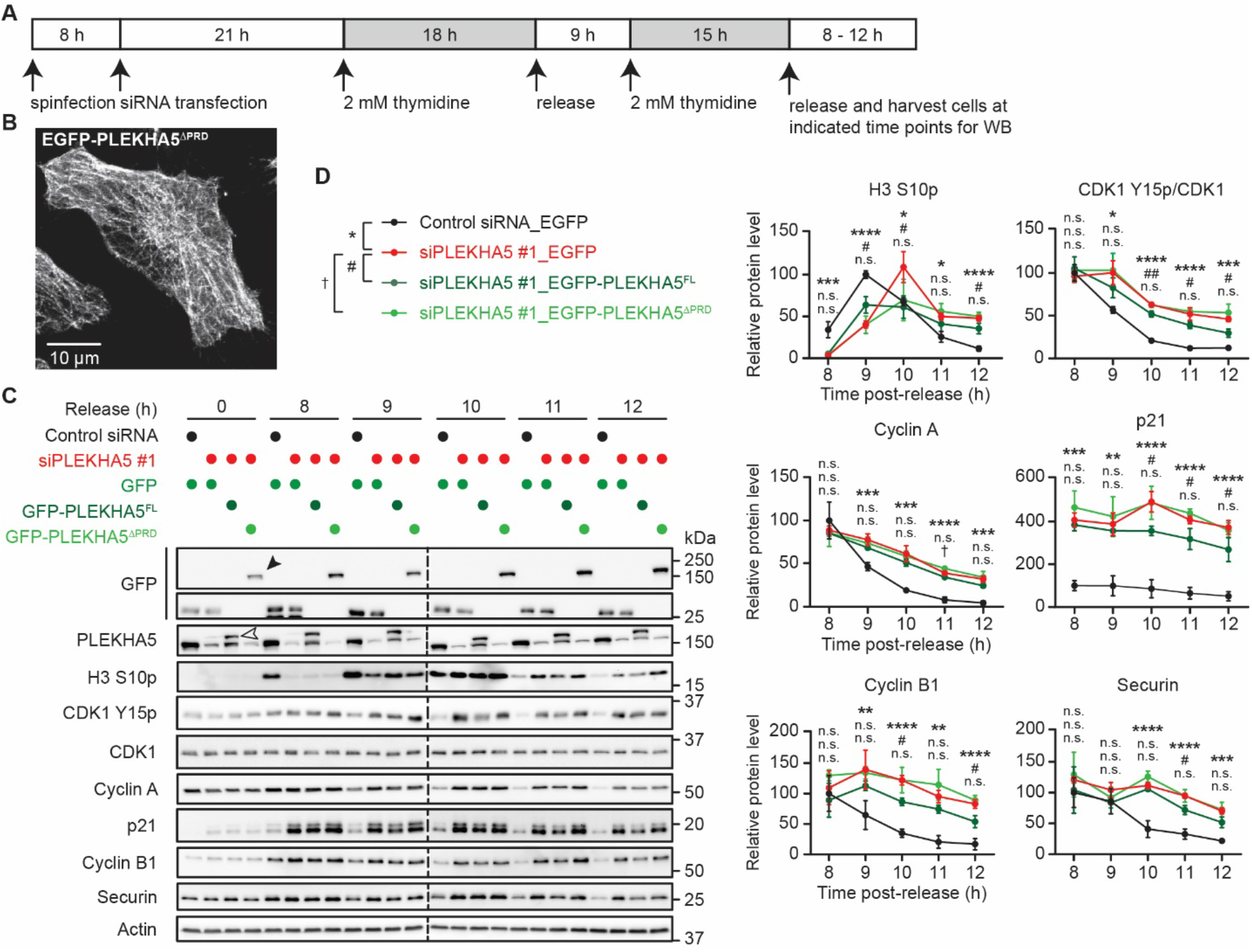
The PLEKHA5–APC/C interaction is required to rescue the accumulation of mitosis markers and APC/C substrates induced by PLEKHA5 knockdown. (A) Schematic representation of experimental timeline. HeLa cells were transduced with lentivirus encoding EGFP, siPLEKHA5-resistant EGFP-PLEKHA5^FL^, or an siPLEKAH5-resistant construct lacking PRD (EGFP-PLEKHA5^δPRD^) via spinfection. Eight h after the spinfection, cells were transfected with PLEKHA5-targeting siRNA #1 or a control siRNA and subjected to DTB. Cells were then released from the block and harvested at indicated time points for Western blot analysis. (B) Confocal microscopy analysis of EGFP-PLEKHA5^δPRD^, showing its microtubule localization. (C–D) Accumulation of mitosis markers and APC/C substrates in siPLEKHA5 samples is partially rescued by exogenous expression of EGFP-PLEKHA5^FL^ but not EGFP-PLEKHA5 ^ΔPRD^. Shown are representative Western blots (C) and quantification of protein levels (D) in HeLa cells transduced with conditioned media containing lentivirus encoding EGFP, siRNA-resistant EGFP-PLEKHA5^FL^, and siRNA-resistant EGFP-PLEKHA5^δPRD^ (n=3). Notes: EGFP-PLEKHA5^FL^ expression is verified by PLEKHA5 blot (hollow arrowhead; this construct is expressed at levels too low for detection by GFP antibody); EGFP-PLEKHA5^ΔPRD^ expression verified by GFP blot (solid arrowhead; this construct does not contain epitope recognized by the PLEKHA5 antibody). ANOVA (one-way, Tukey): n.s. not significant; *, #, and † p < 0.05; **, ##, and †† p < 0.01; ***, ###, and ††† p<0.001; ****, ####, and †††† p < 0.0001.

### PLEKHA5 knockdown attenuates mitotic progression

Though the above data using DTB synchronization imply the importance of PLEKHA5 in mitotic regulation, it is possible that effects of PLEKHA5 knockdown in M phase could result from general effects on cell cycle position. We therefore took additional approaches to assess roles of PLEKHA5 knockdown in M phase progression.

First, we examined the levels of the same G2/M markers and APC/C substrates after a DTB and release but in the presence of the phosphodeficient CDK1 T14A/Y15F (AF) mutant (**Figure S3A**). This CDK1 AF mutant, which cannot possess inhibitory phosphorylation marks, is constitutively active, and its expression induces cells to partially bypass G2/M checkpoint (Jin et al., 1996; Norbury et al., 1991; Ayeni et al., 2014; Szmyd et al., 2019). Compared to control, cells expressing CDK1(AF)-3xFLAG with control siRNA treatment showed increased mitosis progression, signified by earlier timepoints for H3 S10p peak expression, a faster decrease in its levels, and reduced accumulation of APC/C substrates (**Figures S3B–C**, compare grey and black line in quantification plots, significance marked by *). These results confirm that expressing CDK1(AF) mutant indeed permits cells to partially bypass the G2/M checkpoint and promote mitosis entry in our system. We then compared the effect of PLEKHA5 knockdown versus control siRNA in CDK1(AF)-expressing cells. Again, we observed a delay in H3 S10p disappearance and elevated levels of endogenous CDK1 Y15p and APC/C substrates upon PLEKHA5 knockdown (**Figures S3B–C**, compare black and red line in quantification plots, significance marked by #), supporting a role for PLEKHA5 in mitosis.

Second, to eliminate negative effects attributable to synchronization methods, we performed temperature- and CO_2_-controlled live-cell imaging over extended time periods of asynchronous cells expressing fluorescent markers of mitosis progression. We imaged cells stably expressing histone H2B-mCherry every 4 min and quantified the time from nuclear envelope breakdown (NEBD) to anaphase onset, signified by chromosomal segregation (**Figure S4A**; Hayward et al., 2019). PLEKHA5 knockdown caused a minor but significant increase in time for cells to enter anaphase (**Figures S4B–C**) but did not exert deleterious effects on mitotic exit, which was measured as time from anaphase to chromosome decondensation (**Figure S4D**). The modest nature of this delay suggests an effect of PLEKHA5 on only a subset of APC/C complexes. Nevertheless, we note that the effect on mitosis in unperturbed, asynchronous cells seen here is similar in magnitude to depletion of the protein phosphatase PP2A-B55, a non-essential but important regulator of mitosis (Hayward et al., 2019). Collectively, these experiments demonstrate that siPLEKHA5 exhibits specific delays in mitotic progression.

### PLEKHA5 promotes APC/C association with CDC20 and activation in prometaphase

Based on the above data implicating PLEKHA5 as an APC/C interactor and modulator of mitotic progression, we hypothesized that PLEKHA5 might influence the assembly of active APC/C and thus affects its catalytic activity. To test this hypothesis, we first examined the composition of APC/C in mitosis. We isolated endogenous APC/C via anti-ANAPC3 IP from HeLa cells that were either asynchronous or synchronized to prometaphase by a 16 h treatment of S-trityl-L-cysteine (STLC) (Skoufias et al., 2006), and we confirmed that several representative structural subunits and its key mitotic co-activator CDC20 were enriched from the cell lysates (**Figure S5A**). We then compared the APC/C enriched from prometaphase cells that had been transfected with either siPLEKHA5 or control siRNA (**Figure 4A**). PLEKHA5 knockdown did not affect assembly of the structural components of APC/C, but the association of APC/C with CDC20 was significantly reduced though the overall level of CDC20 was not affected by siPLEKHA5 treatment (**Figures 4B–C** and **S5B–C**). To measure the effect of this decreased CDC20 association on APC/C function, we assessed the E3 ligase activity of APC/C^CDC20^ using in vitro ubiquitination assays (Oh et al., 2020). We reconstituted the ubiquitination cascade using the enriched APC/C and other required components, including E1 (UBE1), E2 (UBE2S and UBE2C/UbcH10), ubiquitin, ATP, and recombinantly purified securin as the APC/C substrate. Western blot analysis of the reaction mixtures revealed that the in vitro ubiquitination of securin was dependent upon the presence of all components, and the APC/C purified by IP was indeed active (**Figure S5D**). Next, we evaluated the catalytic activities of APC/C^CDC20^ purified from control or siPLEKHA5 cells synchronized to prometaphase. Upon PLEKHA5 knockdown, the in vitro polyubiquitination of securin by APC/C was significantly decreased, which was consistent with a decrease in APC/C loading with CDC20 caused by siPLEKHA5 (**Figure 4D–E**). These results indicate that depletion of PLEKHA5 from prometaphase cells antagonizes the loading of CDC20 onto APC/C and thus impairs the catalytic activity of the E3 ubiquitin ligase towards its substrates.

**Figure 4.**
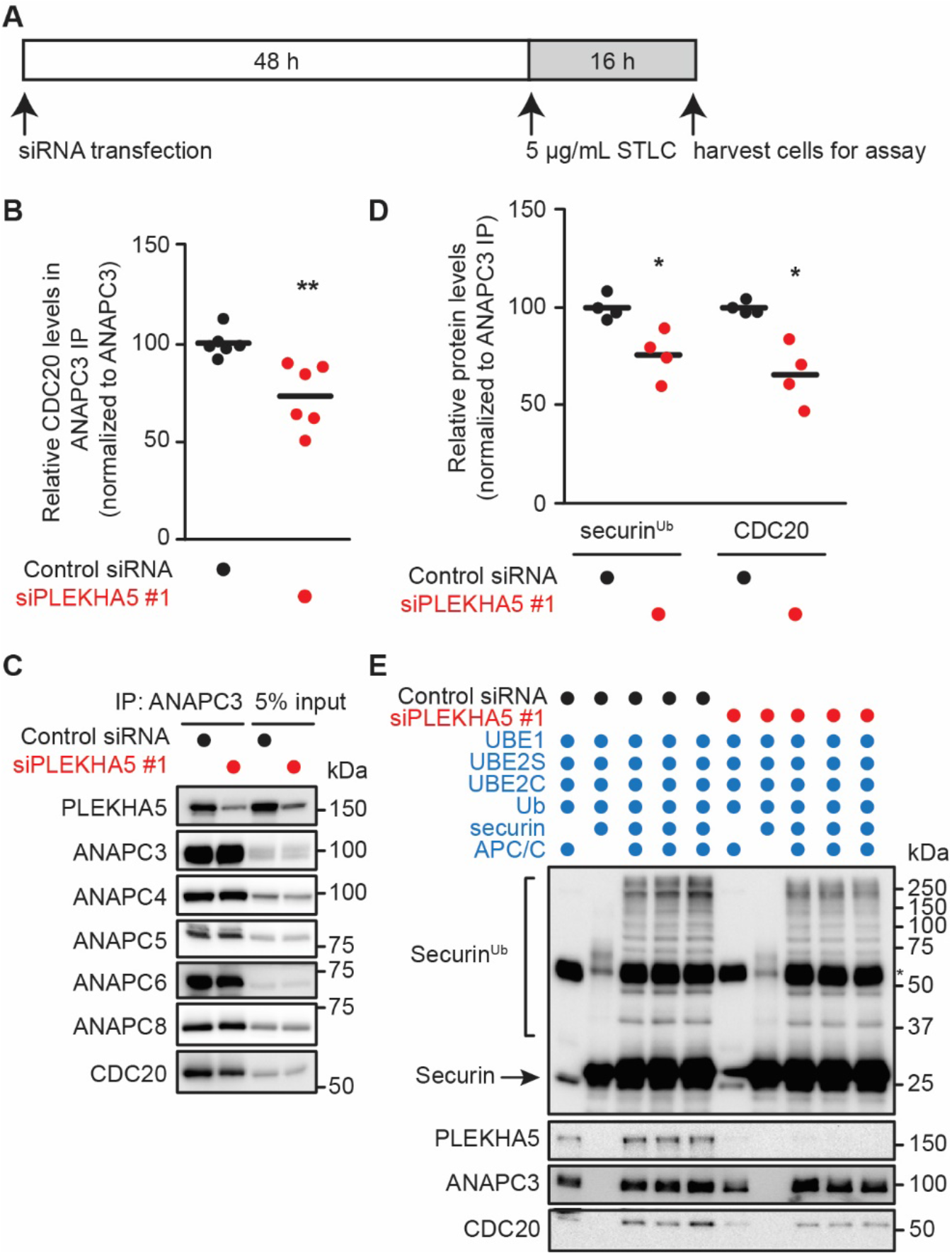
PLEKHA5 knockdown reduces CDC20 loading of APC/C in prometaphase and diminishes the catalytic activity of APC/C. (A) Schematic representation of experimental timeline. HeLa cells were transfected with PLEKHA5-targeting siRNA or a control siRNA. After 48 h, cells were synchronized to prometaphase by STLC treatment. Prometaphase cells were detached from the plate by gentle tapping and collected for use in assays. (B–C) siPLEKHA5 #1 treatment causes a decrease in APC/C association with CDC20. Shown are representative Western blot analysis of co-IP experiments for endogenous ANAPC3 (C) and quantification of CDC20 levels in IP samples (B) (n=6). (D–E) In vitro ubiquitination assays indicate that PLEKHA5 siRNA causes a reduction in APC/C E3 ligase activity. APC/C was purified from prometaphase-synchronized HeLa cells treated with control or PLEKHA5 siRNA #1 using α-ANAPC3 IP. In vitro ubiquitination reactions with the APC/C substrate securin were set up by addition of all of the indicated components shown in (E). Reactions were incubated at 30 °C for 30 min with gentle agitation, followed by denaturation and Western blot, probing for securin. Shown are representative Western blots for the assay (E) and quantification of polyubiquitinated securin (securin^Ub^) and CDC20 levels in the reactions (D). IgG heavy chain from the APC/C purification is marked by an asterisk, and complete reactions (lanes 3–5 and 8–10) are shown in technical triplicate for each biological replicate (n=4 biological replicates plotted in D). Student’s t-test: * p < 0.05; ** p < 0.01.

### Development of a proximity biotinylation tool to probe the microtubule localization of endogenous PLEKHA5, APC/C, and its substrates

Given that PLEKHA5 is a microtubule-localized protein (**Figures 1B–C**; Sluysmans et al., 2021a), that the PLEKHA5–APC/C interaction was critical to ensure proper mitotic progression (**Figure 3C**), and that the presence of PLEKHA5 enables APC/C to be properly charged with CDC20 and fully active in mitosis (**Figure 4E**), we hypothesized that the effects of PLEKHA5 on APC/C function could be due to a role in recruiting this E3 ligase to the microtubule network. To test this hypothesis, we needed to examine the co-localizations of PLEKHA5, APC/C, CDC20 and the substrates on microtubules. Previous studies using immunofluorescence have reported multiple localizations of APC/C, including the cytosol, the prophase nucleus, and, critically, numerous parts of the microtubule network (Sivakumar and Gorbsky, 2015). However, commercial antibodies for APC/C have proven unreliable for immunofluorescence, and GFP fusions of APC/C components have failed to properly assemble into functional APC/C complexes (Huang and Raff, 2002; Acquaviva et al., 2004). A recent high-throughput effort (OpenCell) has generated CRISPR knockin cell lines with GFP fusions to >1300 human proteins expressed endogenously (Cho et al., 2022), yet for several APC/C subunits in this collection, no reliable localization was observed to intracellular organelles or structures.

The deficiencies of these conventional tools motivated us to develop an alternative strategy harnessing TurboID-based proximity labeling (Samavarchi-Tehrani et al., 2020; Branon et al., 2018; Qin et al., 2021) to ascertain the microtubule localization of endogenous PLEKHA5, APC/C and other proteins (**Figure 5A**). Proximity labeling-based tools exist for tagging specific subsets of the microtubule network, e.g., the centrosome–cilium interface, non-centrosomal microtubule organizing centers, kinetochores, and dynein-associated proteins (Gupta et al., 2015; Redwine et al., 2017; Remnant et al., 2019; Sanchez et al., 2021). To our knowledge, however, tools for broad labeling of proteins associated with the microtubule network in mammalian cells were not available. Therefore, we developed two microtubule-targeted TurboID (MT-TurboID) constructs by fusing V5-tagged TurboID to two microtubule-associated proteins: doublecortin (DCX) and the microtubule-binding domain of ensconsin/MAP7 (EMTB) (Ettinger et al., 2016; Miller and Bement, 2008).

**Figure 5.**
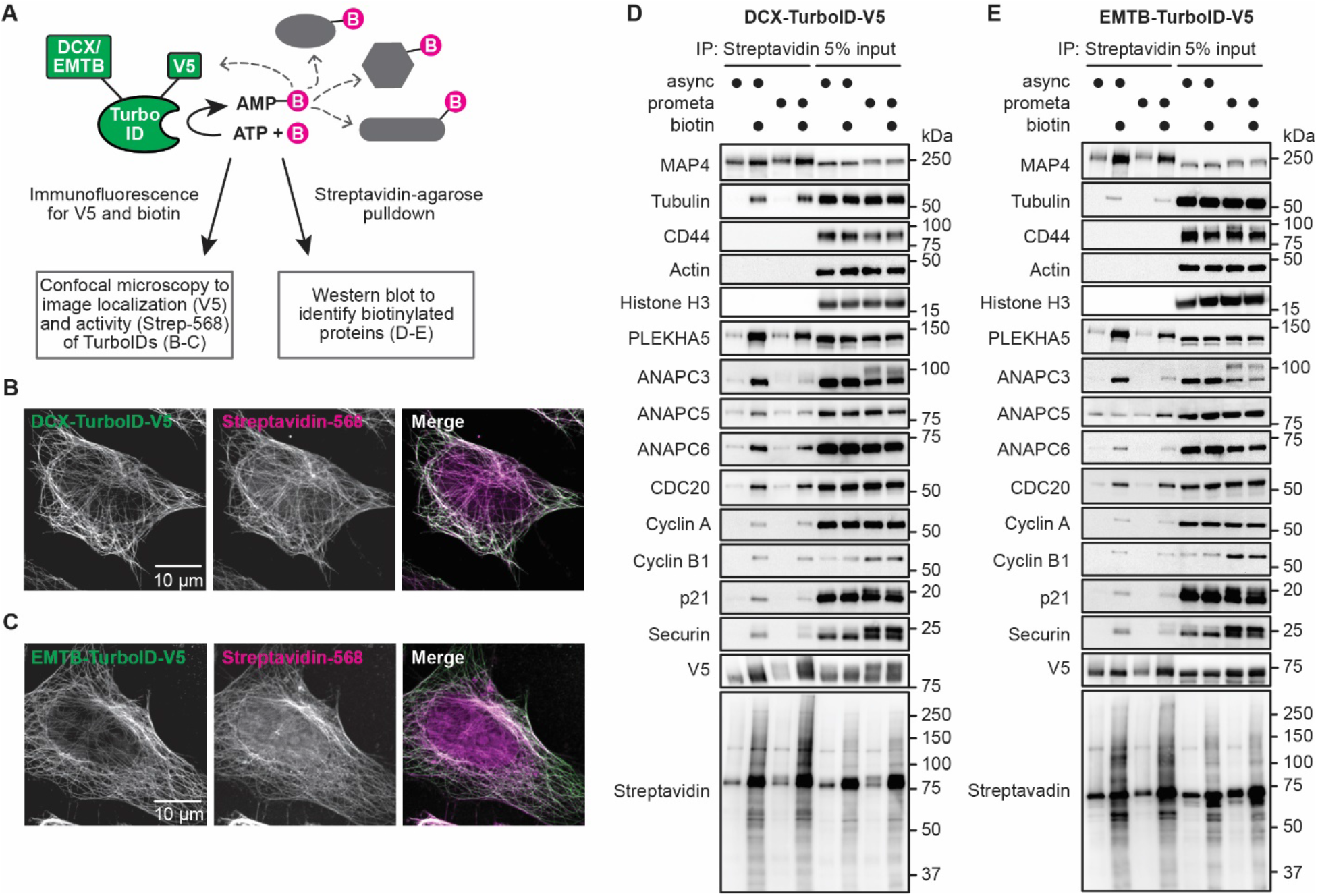
Proximity biotinylation using microtubule-targeted TurboIDs reveals localization of endogenous PLEKHA5 and APC/C to the microtubule network. (A) Schematic representation of proximity biotinylation using MT-TurboID for visualization and identification of microtubule-associated proteins. Two MT-TurboID constructs were generated, which contained fusions of V5-tagged TurboID to one of two microtubule-localizing proteins: doublecortin (DCX) and the microtubule-binding domain of ensconsin/MAP7 (EMTB). (B–C) Both microtubule-targeting TurboID constructs show correct cellular localization and are active. HeLa cells were transfected with DCX-TurboID-V5 (B) or EMTB-Turbo-V5 (C) for 30 h and were then treated with 500 μM biotin for 10 min. Cells were fixed, stained with α-V5 (green) and streptavidin-568 (magenta), and imaged by confocal microscopy. (D–E) PLEKHA5, APC/C subunits, and APC/C substrates, but not markers of other subcellular locations, were biotinylated by microtubule-targeting TurboID constructs in both asynchronous and prometaphase-synchronized HeLa cells. HeLa cells were transfected with DCX-TurboID-V5 (D) or EMTB-Turbo-V5 (E) for 30 h and were either left to be asynchronous or synchronized to prometaphase using STLC (5 μg /mL for 16 h). Biotin was then added to cells for 10 min before cell lysates were harvested. Lysates were subjected to streptavidin-agarose enrichment of biotinylated proteins, and Western blot was performed to identify biotinylated proteins.

We first validated the localization and biotinylation activity of the two MT-TurboID constructs, DCX-TurboID-V5 and EMTB-TurboID-V5, by performing IF against the V5 tag and biotin (**Figures 5B–C**). We then validated the specificity of the MT-TurboIDs using streptavidin-agarose enrichment of biotinylated proteins followed by Western blot analysis (**Figures 5D–E**). Both MT-TurboID constructs successfully labeled Microtubule Associated Protein 4 (MAP4), α-tubulin, and PLEKHA5 upon the addition of exogenous biotin but showed negligible biotinylation of markers of the plasma membrane (CD44), the actin cytoskeleton (actin), and the nucleus (histone H3), demonstrating that the activity of MT-TurboIDs was selectively confined to proteins on microtubules. Importantly, in the same experiment, representative APC/C subunits ANAPC3, ANAPC5 and ANAPC6, the co-activator CDC20, and APC/C substrates including cyclin A, cyclin B1, p21, and securin were biotinylated by both MT-TurboIDs in either asynchronous or prometaphase-synchronized cells (**Figure 5D–E**). These results reveal that APC/C structural subunits, CDC20, and several APC/C substrates all exhibit microtubule localization both in interphase and in M phase and further validate MT-TurboIDs as general ‘in vivo biochemistry’ tools for interrogating the microtubule localization of target proteins of interest.

### PLEKHA5 depletion caused a decrease in the microtubule localization of APC/C subunits, but not CDC20

Finally, to elucidate the requirement for PLEKHA5 in promoting APC/C localization to microtubules, we evaluated the extent of microtubule association of APC/C in control vs. siPLEKHA5 cells using MT-TurboID proximity biotinylation, enrichment of biotinylated proteins, and Western blot (**Figure 6A**). We confirmed that treating cells with siPLEKHA5 did not alter the expression level nor the activity of MT-Turbos (**Figures 6C and E**, V5 and streptavidin blots). Upon knockdown of PLEKHA5 in either asynchronous populations or cells synchronized to prometaphase, several representative subunits of the APC/C complex exhibited a significant decrease in microtubule localization marked by both MT-TurboID constructs (**Figure 6B–E**). However, in these same experiments, there was no significant change in levels of microtubule-associated CDC20 upon PLEKHA5 knockdown (**Figures 6B–E**). These data suggest that PLEKHA5 plays a role in recruitment of a pool of APC/C complexes to microtubules and are consistent with a model wherein PLEKHA5 functions as an APC/C adaptor to promote its localization to the microtubule and thus enhance the efficiency of CDC20 loading onto this pool of APC/C to promote ubiquitination of microtubule-localized substrates during mitosis (**Figure 7**).

**Figure 6.**
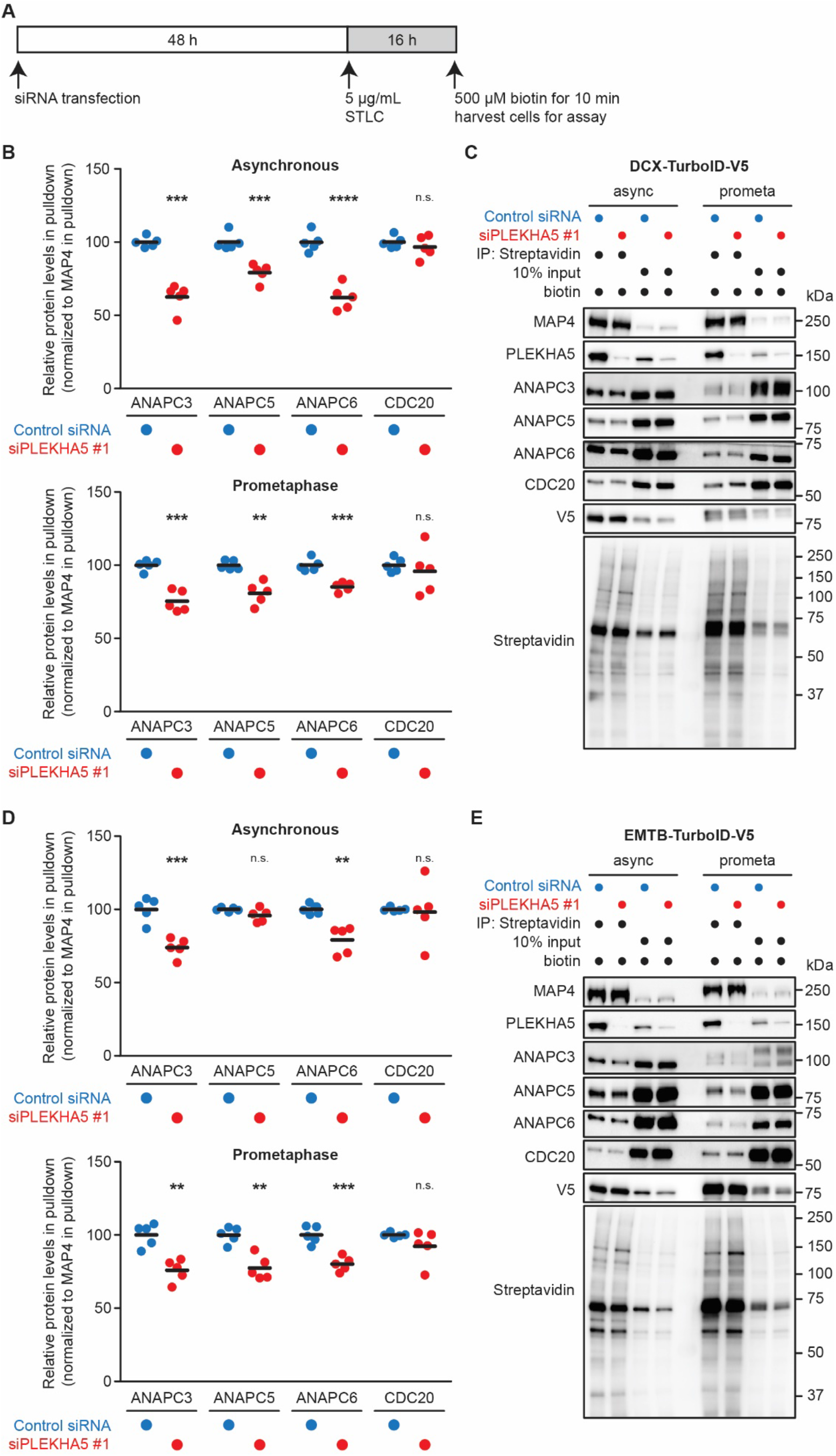
PLEKHA5 knockdown causes decrease in association of APC/C, but not CDC20, with microtubules. (A) Schematic representation of experimental timeline. HeLa cells stably expressing DCX-Turbo-V5 or EMTB-Turbo-V5 (see Figure 7 legend) were subjected to siPLEKHA5 or a control siRNA. 48 h post siRNA transfection, cells were either left asynchronous or synchronized to prometaphase by treatment with STLC (5 μg/mL for 16 h), followed by biotinylation, lysis, enrichment using streptavidin-agarose, Western blot analysis. (B–E) Knockdown of PLEKHA5 leads to reduced levels of APC/C subunits in streptavidin pulldowns from HeLa cells stably expressing DCX-TurboID-V5 (B–C) or EMTB-TurboID-V5 (D–E). Shown are representative blots for the assay (C and E) and quantifications of protein levels (B and D) in streptavidin pulldowns (n=5). Student’s t-test: n.s. not significant; ** p < 0.01; *** p<0.001; **** p < 0.0001.

**Figure 7.**
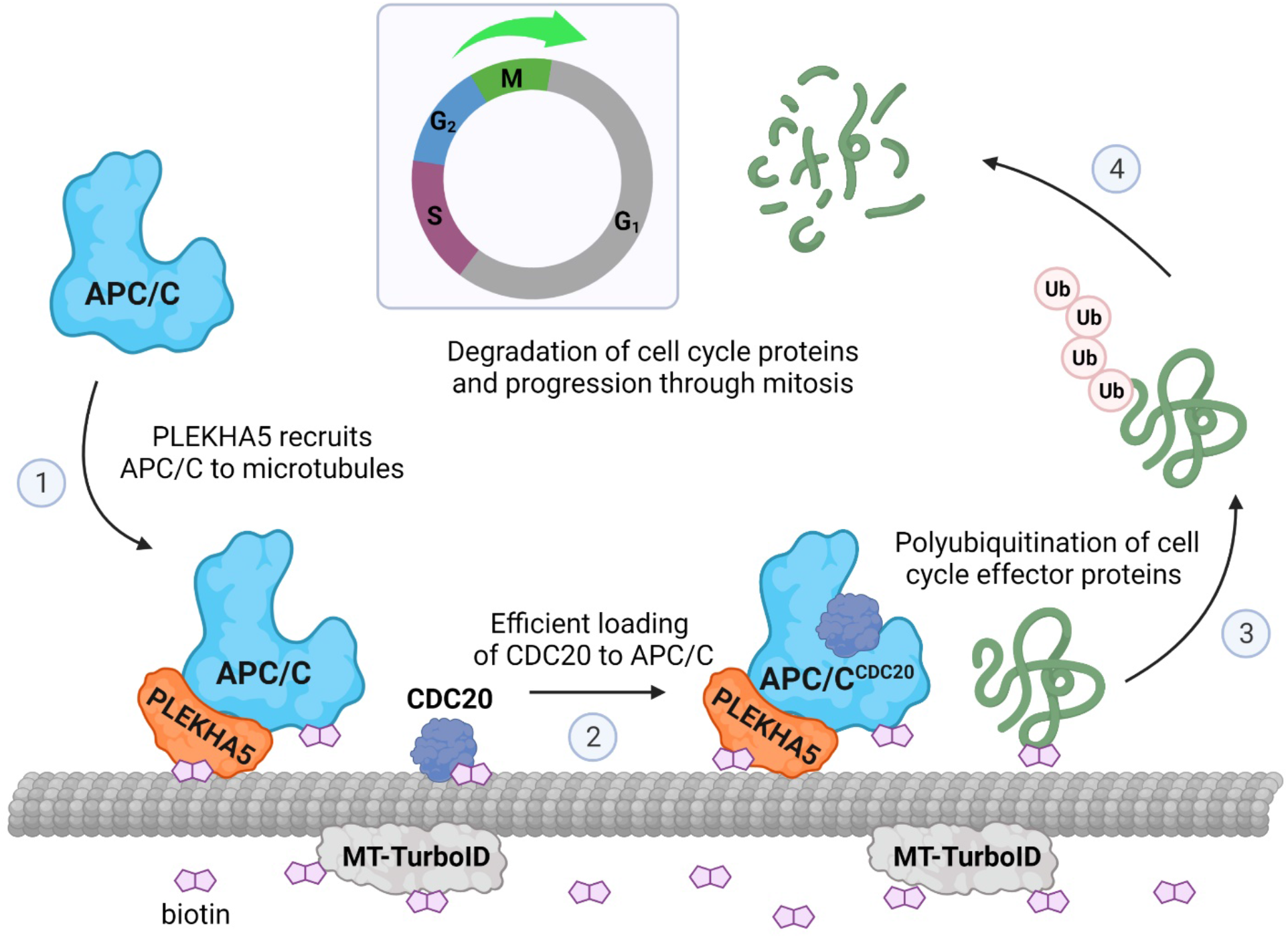
Model for PLEKHA5 function in mitosis as a microtubule-localized adaptor that regulates spatial organization of the APC/C E3 ubiquitin ligase. PLEKHA5 is a microtubule-resident protein that recruits a pool of APC/C to the microtubule cytoskeleton, as demonstrated by its biotinylation by microtubule-localized proximity labeling tools (MT-TurboID), thus increasing the local concentration of APC/C along the microtubule network. Upon mitotic phosphorylation, this pool of APC/C can efficiently interact with its co-activator CDC20 and subsequently polyubiquitinate its mitotic substrates. In this manner, PLEKHA5 organizes APC/C in space to facilitate the timely turnover of key mitotic proteins to promote progression through mitosis (Figure created with BioRender).

## DISCUSSION

PLEKHA5 is a member of the PLEKHA4–7 protein family, several members of which are linked to cancer or cancer-related cellular processes such as proliferation or migration. PLEKHA4 levels are elevated in melanoma compared to healthy melanocytes, and depletion of PLEKHA4 attenuates melanoma cell proliferation in vitro and tumor xenograft growth in vivo (Shami Shah et al., 2021). PLEKHA7 is upregulated in colorectal cancer and supports the growth, adhesion, and invasion of KRAS-mutant colorectal cancer cells (Jeung et al., 2021). PLEKHA5 itself is critical for proliferation and invasiveness of melanoma cells with a high propensity for brain metastasis, supported by both in vitro migration assays that model blood-brain-barrier penetration and clinical data showing positive correlations between PLEKHA5 levels and early onset of brain metastasis (Jilaveanu et al., 2015). Other studies have implicated PLEKHA5 in regulation of malignant progression in diffuse-type gastric carcinoma and breast cancer metastasis (Nagamura et al., 2021; Liu et al., 2020). These reports underscore the importance of elucidating the fundamental molecular and cellular functions of PLEKHA5, particularly those that impact regulation of the cell cycle. Though cancer cell survival is not absolutely dependent upon PLEKHA5 (DepMap and Broad, 2019), non-essential genes can still function as important regulators of the cell cycle (Hayward et al., 2019), and understanding their functional mechanisms can reveal important insights into cell cycle regulation.

We found that PLEKHA5 exhibits a microtubule localization in HeLa cells mediated by its N-terminal tandem WW domains, consistent with a recent study of the WW-PLEKHA proteins (Sluysmans et al., 2021a). We identified a protein-protein interaction between PLEKHA5 and the E3 ubiquitin ligase APC/C, to which PLEKHA5 binds via its unstructured proline-rich domain. We reasoned that PLEKHA5 is likely not a substrate for APC/C during mitosis, because PLEKHA5 levels remained stable throughout G2 and M phases and it does not possess APC/C destruction motifs such as the D-box (Glotzer et al., 1991), KEN-box (Pfleger and Kirschner, 2000), or ABBA motif (DiFiore et al., 2015) within its primary amino acid sequence. Instead, we hypothesized that PLEKHA5 might be a microtubule-localized APC/C adaptor.

Multiple lines of evidence support the idea that PLEKHA5 functions to ensure advancement of mitosis and proper turnover of APC/C substrates. Depletion of PLEKHA5 by siRNA antagonized mitotic progression and led to accumulations of APC/C substrates in a manner dependent upon the PLEKHA5–APC/C interaction. To achieve precise spatiotemporal control over substrate ubiquitination and degradation, APC/C activity is subjected to multiple layers of regulation, including interaction with E2 enzymes (Rodrigo-Brenni and Morgan, 2007; Yau et al., 2017; Brown et al., 2016), binding to inhibitors (Reimann et al., 2001; Machida and Dutta, 2007; Alfieri et al., 2016; Yamaguchi et al., 2016) and co-activators (Visintin et al., 1997; Kimata et al., 2008), and phosphorylation (Fujimitsu et al., 2016; Qiao et al., 2016). In early mitosis, CDK1 and polo-like kinase 1 (PLK1) phosphorylate the two APC/C subunits, ANAPC3 and ANAPC1, within their disordered loop domains, and consequently the autoinhibitory loop of ANAPC1 is dislodged, exposing a CDC20 binding site and thus allowing stable interaction of this key mitotic co-activator with APC/C (Zhang et al., 2016).

We did not observe differences in the phosphorylation patterns of ANAPC3 in prometaphase-synchronized control vs. siPLEKHA5 cells, indicating that PLEKHA5 does not affect APC/C phosphorylation status. However, APC/C complexes isolated from prometaphase-synchronized cells depleted of PLEKHA5 exhibited a reduced interaction with CDC20 and subsequent decrease in catalytic activity towards a mitotic substrate, securin, in vitro. Because the presence or absence of PLEKHA5 did not affect the APC/C phosphorylation status, APC/C in both control and siPLEKHA5 cells should have a similar affinity toward CDC20, whose overall levels did not change as a result of siPLEKHA5 treatment. Therefore, we hypothesized that, rather than tuning the affinity of co-activator binding, PLEKHA5 impacts CDC20-depedent APC/C activation through spatial regulation, that is by bringing APC/C to a particular subcellular localization, namely the microtubule cytoskeleton, where the APC/C could more efficiently access CDC20.

In contrast to the temporal control of APC/C-mediated polyubiquitination, which has been extensively studied (Pines, 2011; Alfieri et al., 2017; Watson et al., 2018), the factors and mechanisms that control the dynamic localizations of APC/C remain elusive (Sivakumar and Gorbsky, 2015), due in part to difficulties in determining APC/C localizations using conventional imaging-based methods such as IF and (even endogenous) fluorescent protein tagging (Huang and Raff, 2002; Acquaviva et al., 2004; Cho et al., 2022). To overcome these challenges and to test the hypothesis that PLEKHA5 mediates the spatial organization of APC/C, we developed a proximity biotinylation approach using microtubule targeted TurboID constructs to assess endogenous protein localizations to the microtubule cytoskeleton. Indeed, we detected biotinylation of PLEKHA5, APC/C, CDC20 and relevant APC/C substrates, confirming that these proteins localize to or very near to microtubules and demonstrating the utility of these MT-TurboIDs as versatile “in vivo biochemistry” tools for examining the microtubule localization of endogenous proteins of interest. The two MT-TurboIDs (DCX-TurboID and EMTB-TurboID) appear to decorate the entire microtubule network rather than specific locations, such as the centrosomes/spindle poles and kinetochores, to which APC/C can localize (Tugendreich et al., 1995; Tischer et al., 2021; Topper et al., 2002; Jörgensen et al., 1998). Additional TurboID constructs localized to these specific substructures within the microtubule network (Gupta et al., 2015; Redwine et al., 2017; Remnant et al., 2019; Sanchez et al., 2021), and different types of microtubule fibers including kinetochore fibers, non-kinetochore fibers, and astral fibers, may in the future be useful for more granular sampling of APC/C localizations and APC/C composition and phosphorylation status at such locations which may be critical for modulating APC/C activity and functions in mitosis (Sivakumar et al., 2014; Torres et al., 2010).

Collectively, this study reveals a new framework for the spatial regulation of APC/C in mitosis. In our model, PLEKHA5 interacts with and recruits a pool of APC/C to the microtubule cytoskeleton. This interaction would increase the local concentration of APC/C along the network and effectively reduce the dimensionality of its search for co-activator and substrates from three dimensions (in the cytosol) to one dimension (along microtubule tracks). Upon mitotic phosphorylation of its subunits and relief of the spindle assembly checkpoint, this pool of APC/C would efficiently encounter its co-activator CDC20 and subsequently be capable of ubiquitinating key mitotic effector proteins. Our study not only reveals a mechanism controlling the spatial regulation of the major E3 ubiquitin ligase in mitosis but also provides a means for understanding how such spatial regulation could be intimately linked to the established temporal regulation of this enzyme that is critical for progression through mitosis. Given the localization of PLEKHA5 and the large number of PLEKHA5-interacting proteins identified in our proteomics experiments, it is possible that PLEKHA5 acts more generally as an adaptor to facilitate protein recruitment to the microtubule cytoskeleton in both interphase and mitosis. Further, our studies provide a mechanism that accounts for a role for PLEKHA5 in promoting progression through mitosis that may be operational in settings where elevated PLEKHA5 levels have been linked to cancer progression. Finally, the MT-TurboIDs developed herein have the potential to serve as broadly useful tools for interrogating the dynamic association of diverse proteins with the microtubule cytoskeleton in many physiological contexts.

## STAR METHODS

### Antibodies

Mouse anti-PLEKHA5, Santa Cruz Biotechnology (sc-390311, RRID: N/A); mouse anti-ANAPC3, Santa Cruz Biotechnology (sc-9972, RRID: AB_627228); rabbit anti-ANAPC4, Bethyl Laboratories (A301-176A, RRID: AB_2227071); rabbit anti-ANAPC5, ABclonal (A7109, RRID: AB_2767664); mouse anti-ANAPC6, Santa Cruz Biotechnology (sc-376091, RRID: AB_10988885); rabbit anti-ANAPC8, Cell Signaling Technology (15100, RRID: AB_2798708); mouse IgG2b isotype control, Cell Signaling Technology (53484, RRID: AB_2799435); normal rabbit IgG control, Cell Signaling Technology (2729, RRID: AB_1031062); mouse anti-GFP, Santa Cruz Biotechnology (sc-9996, RRID: AB_627695); mouse anti-phospho-Histone H3 (Ser10), Cell Signaling Technology (9706, RRID: AB_331748); rabbit anti-phospho-CDK1 (Tyr15), Cell Signaling Technology (9111, RRID: AB_331460); mouse anti-CDK1, Santa Cruz Biotechnology (sc-54, RRID: AB_627224); mouse anti-cyclin A, Santa Cruz Biotechnology (sc-239, RRID: AB_627334); mouse anti-cyclin B1, Santa Cruz Biotechnology (sc-245, RRID: AB_627338); rabbit anti-p21, Cell Signaling Technology (2947, RRID: AB_823586); mouse anti-securin, Santa Cruz Biotechnology (sc-56207, RRID: AB_785382); mouse anti-actin, MP Bio (08691001, RRID: AB_2335127); mouse anti-GAPDH, GeneTex (GTX78213, RRID: AB_625368); rabbit anti-FLAG, Sigma-Aldrich (F7425, RRID: AB_439687); rabbit anti-CDC20, Cell Signaling Technology (4823, RRID: AB_10549074); mouse anti-CDC20, Santa Cruz Biotechnology (sc-13162, RRID: AB_628089); mouse anti-MAP4, Santa Cruz Biotechnology (sc-390286, RRID: N/A); mouse anti-α tubulin, Sigma-Aldrich (T5168, RRID: AB_477579); mouse anti-CD44, Cell Signaling Technology (3570, RRID: AB_2076465); rabbit anti-Histone H3, Cell Signaling Technology (4499, RRID: AB_10544537); mouse anti-V5, Bio-Rad (MCA1360GA, RRID: AB_567249); goat anti-mouse HRP, Jackson ImmunoResearch Labs (115-035-146, RRID: AB_2307392); goat anti-rabbit HRP, Jackson ImmunoResearch Labs (111-035-144, RRID: AB_2307391); donkey anti-mouse Alexa Fluor 488, Invitrogen (A-21202, RRID: AB_141607).

### Chemicals and recombinant proteins

Lipofectamine 2000, Invitrogen (11668019); Lipofectamine RNAiMAX, Invitrogen (13778150); Polybrene Infection/Transfection Reagent, Sigma-Aldrich (TR-1003); GFP-Trap magnetic agarose, ChromoTek (gtma-20); Trypsin Gold Mass Spectrometry Grade, Promega (V5280); Protein G Sepharose, BioVision (6511); Thymidine, Chem-Impex (00306); Propidium iodide, Sigma-Aldrich (P4170); RNase A, Research Products International (R21750); Nocodazole, AdipoGen (AG-CR1-0019); (+)-S-Trityl-L-cysteine, Alfa Aesar (L14384); Puromycin dihydrochloride, Sigma-Aldrich (P8833); Blasticidin S hydrochloride, 10 mg/ml in HEPES buffer, Alfa Aesar (J67216); Hygromycin B, 50 mg/mL in PBS, Invitrogen (10687010); cOmplete Protease Inhibitor Cocktail, Roche (5056489001); Sodium β-glycerophosphate pentahydrate, Alfa Aesar (L03425); Sodium fluoride, Chem-Impex (01523); Sodium pyrophosphate decahydrate, Fisher (S390); Sodium orthovanadate, MP Bio (0215966410); UBE1, R&D Systems (E-304); UBE2C, R&D Systems (E2-654); UBE2S, R&D Systems (E2-690); securin, Abcam (ab87664); D-biotin, Chem-Impex (00033); Streptavidin Alexa Fluor 568, Invitrogen (S11226); Pierce™ High Capacity Streptavidin Agarose, Thermo Fisher (20359); Streptavidin-conjugated HRP, GeneTex (GTX85912); ProLong™ Diamond Antifade Mountant with DAPI, Thermo Fisher (P36971); Clarity™ Western ECL Substrate, Bio-Rad (1705061).

### siRNA duplexes

Negative control siRNA

Sense (RNA): 5’-rCrGrUrUrArArUrCrGrCrGrUrArUrArArUrArCrGrCrGrUAT-3’
Antisense (RNA): 5’-rArUrArCrGrCrGrUrArUrUrArUrArCrGrCrGrArUrUrArArCrGrArC-3’

siPLEKHA5 #1

Sense: 5’ -rArGrArCrCrUrGrArArGrArArGrUrArGrArUrArUrUrGrATG-3’
Antisense: 5’-rCrArUrCrArArUrArUrCrUrArCrUrUrCrUrUrCrArGrGrUrCrUrGrU-3’

siPLEKHA5 #2

Sense: 5’ -rGrUrUrGrCrArArCrCrArUrGrArCrArUrCrUrGrArArGrAAA-3’
Antisense: 5’ -rUrUrUrCrUrUrCrArGrArUrGrUrCrArUrGrGrUrUrGrCrArArCrArG-3’

### Plasmids and cloning

A PLEKHA5 cDNA (Uniprot Q9HAU0, a gift from Haiyuan Yu, Cornell University) was cloned into the pEGFP-C1 vector (Clontech) using BamHI and SalI to generate EGFP-PLEKHA5^FL^. Subsequently, fragments and deletions of PLEKHA5 were generated by standard or overlap PCR, and inserts were cloned to pEGFP-C1 vector, again using BamHI and SalI, and to pEGFP-N1-vector with NheI and SalI. Also, two deletion constructs were generated by QuikChange XL Site-directed mutagenesis kit (Agilent). The amino acid sequences of the fragments and deletions are the following: PLEKHA5^WW^ (1-105), PLEKHA5^H-BP-PH^ (143-271), PLEKHA5^WW-H-BP-PH^ (1-271), PLEKHA5^PRD^ (272-628), PLEKHA5^CC-IDR^ (629-1116), PLEKHA5^ΔWW^ (106-1116), PLEKHA5^δH-BP-PH^ (1-142, 272-1116), PLEKHA5^ΔPRD^ (1-271, 629-1116), PLEKHA5^ΔCC-IDR^ (1-628). For generation of stable HeLa cells used in the proteomics experiment, EGFP-PLEKHA5 was subcloned into the pCDNA5-FRT vector (Thermo Fisher) using NheI and BamHI.

For rescue experiments, silent mutations were incorporated into EGFP-PLEKHA5^FL^ and EGFP-PLEKHA5^δPRD^ to generate siPLEKHA5 #1-resistant constructs using the QuikChange XL Site-directed mutagenesis kit. The following codons within the siRNA targeting region were mutated: R629, E631, E632, V633, I635. The siRNA-resistant EGFP-PLEKHA5^FL^ and EGFP-PLEKHA5^ΔPRD^ were then subcloned to an engineered lentiviral transduction vector pCDH-CMV-MCS using NheI and BamHI. The pCDH-CMV-MCS vector was generated by removing the puromycin resistance expression cassette from the pCDH-CMV-MCS-EF1α-Puro vector (a gift from Jan Lammerding, Cornell University) using NotI and SalI, followed by blunting the sticky ends with T4 DNA polymerase (NEB). The blunted DNA fragment was then phosphorylated by T4 Polynucleotide Kinase (NEB) and re-ligated with T4 DNA Ligase (Thermo Fisher). Furthermore, the NotI restriction digestion site was restored, and an AscI site was added upstream of the XbaI site using the QuikChange XL Site-directed mutagenesis kit to generate the final pCDH-CMV-MCS vector.

To generate the lentiviral plasmid encoding CDK1(AF)-3xFLAG, a CDK1(AF) cDNA (Addgene, 39872) was cloned into p3xFLAG-CMV-14 vector (Sigma-Aldrich) using NotI and KpnI. Subsequently, CDK1(AF)-3xFLAG was inserted into the pCDH-CMV-MCS vector for lentivirus production. EGFP-PCNA was cloned by swapping out mTagBFP2 in mTagBFP2-PCNA (Addgene, 85725) for EGFP (digested from EGFP-C1 vector) using AgeI and BspEI, and then EGFP-PCNA was cloned into the pCDH-CMV-MCS-EF1α-Puro vector by XbaI and NotI to yield the lentiviral plasmid.

For TurboID experiments, mammalian-expression plasmids DCX-TurboID-V5 and EMTB-TurboID-V5 were cloned by replacing the C1(1-29) sequence in C1(1-29)-TurboID-V5-pCDNA3 (Addgene, 107173) with DCX (amplified from pLenti6-DCX-EGFP, Addgene, 80599) and EMTB (amplified from EMTB-mCherry, Addgene, 26742) respectively, using HindIII and NotI. The lentiviral plasmid for EMTB-TurboID-V5 was made by cloning the EMTB-TurboID-V5 cassette into the pCDH-CMV-MCS-EF1α-Puro vector via EcoRI and BamHI. The Lentiviral plasmid for DCX-TurboID-V5 was cloned by inserting the DCX-TurboID-V5 cassette into NotI-digested pCDH-CMV-MCS-EF1α-Puro vector via Gibson assembly.

### Cell culture

Flp-In T-Rex HeLa (Thermo Fisher) and HEK 293TN cells (Anthony Bretscher, Cornell University) were cultured in DMEM (Corning) supplemented with 10% FBS (Corning) and 1% penicillin/streptomycin (P/S, Corning) at 37 °C in a 5% CO_2_ atmosphere. HEK 293TN cells were also supplemented with 1% sodium pyruvate (Corning). Stable expression of GFP-PLEKHA5 was achieved by transfecting Flp-In T-Rex HeLa cells with flippase (pOG44, Thermo Fisher) and pCDNA5-FRT-EGFP-PLEKHA5 following the manufacturer’s protocol (Thermo Fisher). After 36 h, cells were selected with 200 μg/mL hygromycin B (Invitrogen). Stable expression of H2B-mCherry was accomplished by transducing pLenti6-H2B-mCherry plasmid (Addgene, 89766) into Flp-In T-Rex HeLa cells and blasticidin (Alfa Aesar) selection at 2.5 μg/mL. Subsequently, lentiviral transduction of pCDH-CMV-EGFP-PCNA-EF1α-Puro followed by puromycin (Sigma-Aldrich) selection at 2 μg/mL yielded the HeLa cell line stably expressing both H2B-mCherry and EGFP-PCNA. HeLa cells stably expressing DCX-TurboID-V5 and EMTB-TurboID-V5 were generated by lentiviral transduction of pCDH-CMV-DCX-TurboID-V5-EF1α-Puro and pCDH-CMV-EMTB-TurboID-V5-EF1α-Puro respectively into Flp-In T-Rex HeLa cells and puromycin selection. Cell lines were used without further authentication, and mycoplasma testing (MycoSensor PCR assay, Agilent) was performed yearly.

### Transfection of plasmids and siRNAs

Plasmid transfections were performed using Lipofectamine 2000 (Invitrogen) following the manufacturer’s protocol but replacing Opti-MEM with Transfectagro (Corning). Cells were incubated with the transfection mix in Transfectagro with 10% FBS for 7–8 h, then the media was exchanged back to regular growth medium. DsiRNA duplexes were purchased from IDT. siRNA transfections were achieved using Lipofectamine RNAiMAX (Invitrogen) according to the manufacturer’s procedures except using Transfectagro instead of Opti-MEM. Twelve to 16 h after incubation with the transfection mix in Transfectagro with 10% FBS, cells were supplemented with the regular growth medium.

### Confocal microscopy

For live-cell imaging, cells were seeded on 35 mm glass bottom culture dishes (14 mm diameter, #1.5 thickness, Matsunami Glass). For immunofluorescence, cells were seeded on 12 mm cover glass (#1.5 thickness, Fisherbrand) in 12-well plates (Corning). Cells were imaged (live cell) or fixed (immunofluorescence) 24-30 h post transfection. For immunofluorescence analysis of PLEKHA5 and MT-TurboID constructs, cells were extracted in Microtubule Stabilizing Buffer (80 mM PIPES pH 6.8, 1 mM MgCl_2_, 5 mM EGTA pH 6.8, 0.5% Triton-X 100) for 30 s and fixed in methanol prechilled to −20 °C for 3 min. The cells were then rehydrated in wash buffer (0.1% Tween-20 in 1X PBS) for 5 min three times, further permeabilized with 0.5% Triton-X 100 in 1X PBS for 5 min and blocked in 2% BSA and 0.1% Tween-20 in 1X PBS (blocking buffer) for 30 min. Cells were incubated with primary antibody in blocking buffer for 1 h, rinsed with wash buffer three times and treated with secondary antibody in blocking buffer for 1 h. Cells were then rinsed with wash buffer three times, with 1X PBS twice, mounted on slides in ProLong Diamond Antifade with DAPI (Thermo Fisher) and incubated overnight in dark before imaging. All steps were performed at room temperature, and slides were stored at 4 °C for long-term storage. Images were acquired via Zeiss Zen Blue 2.3 software on a Zeiss LSM 800 confocal laser scanning microscope equipped with Plan Apochromat objectives (40x 1.4 NA) and two GaAsP PMT detectors. Solid-state lasers (405, 488, and 561 nm) were used to excite DAPI, EGFP/Alexa Fluor 488 and mCherry/Alexa Fluor 568 respectively. Live-cell time-series movies were acquired using definite focus. Depending on the experiments, images and movies were acquired either at room temperature, or at 37 °C with a 5% CO_2_ atmosphere. Acquired images were analyzed using FIJI.

### Immunoprecipitation and Western blots

Cells were harvested and lysed in IP lysis buffer (150 mM NaCl, 1% NP-40, 0.25% sodium deoxycholate, 5 mM EDTA, 50 mM Tris pH 7.5) supplemented with cOmplete protease inhibitor cocktail (Roche) on ice. Cell lysates were sonicated for 6 pulses at 20% intensity and centrifuged at 13,000 *g* for 5 min at 4 °C. Protein concentration of the clarified cell lysates was quantified using the BCA assay (Thermo Fisher), and a small fraction of the clarified cell lysates was separated and normalized as input. The remaining cell lysates were subjected to immunoprecipitation using GFP-Trap Magnetic Agarose (Chromotek) with rotation at 4 °C overnight (12–16 h). The magnetic resin was then separated by a magnetic tube rack, washed three times with lysis buffer (at least 10 times the bead volume), denatured and analyzed by SDS-PAGE and Western blot with detection by chemiluminescence using Clarity Western ECL substrate (Bio-Rad), or by mass spectrometrybased proteomics analysis, as described in detail in the next section. For immunoprecipitation against endogenous proteins using soluble antibodies, the clarified cell lysates were incubated with primary antibodies (2 μg antibody per mg of input for IgG control, PLEKHA5, ANAPC3 and ANAPC4, and 1:150 dilution for ANAPC8) for 2 h at 4 °C with rotation, followed by addition of Protein G Sepharose (BioVision) and rotation at 4 °C overnight. The resin was then centrifuged for 5 min at 1000 *g*, washed three times with lysis buffer (at least 10 times of the bead volume), denatured and analyzed by SDS-PAGE and Western blot with detection by chemiluminescence using Clarity Western ECL substrate.

### SILAC labeling and mass spectrometry-based proteomics analysis

For quantitative proteomics, Flp-In HeLa cells stably expressing EGFP-PLEKHA5 was cultured in “heavy” SILAC DMEM media (Thermo) supplemented with 10% dialyzed FBS (Corning) and 1% P/S for at least five passages to allow full labeling of cells before they were used in experiments. “Light” EGFP-expressing HeLa cell line was generated previously (Shami Shah et al., 2019). “Heavy” SILAC DMEM contained L-lysine ^13^C_6_, ^15^N_2_ and L-arginine ^13^C_6_, ^15^N_4_. “Light” SILAC DMEM contained ^12^C_6_, ^14^N_2_ and L-arginine ^12^C_6_, ^14^N_4_. Cells were lysed and immunoprecipitated with GFP-Trap Magnetic Agarose as described above and processed for mass spectrometry following published procedures (Liu et al., 2017; Sanford et al., 2021).

The resin was washed four times with lysis buffer and incubated with elution buffer (100 mM Tris pH 8.0, 1% SDS) for 5 min at 95 °C. The supernatant from “light” and “heavy” samples were collected and mixed in a 1:1 ratio. The mixed sample was then reduced with 10 mM DTT (65 °C, 10 min) and alkylated with 25 mM iodoacetamide (in 50 mM Tris pH 8.0, room temperature, 15 min in dark). Proteins from the mixed sample were precipitated in a solution containing 50% acetone, 49.9% ethanol and 0.1% acetic acid (PPT solution) on ice for 1 h and pelleted at 17,000 *g* for 10 min. The protein pellet was washed with PPT solution once, pelleted again, air-dried and resuspended in urea/Tris solution (8 M urea, 50 mM Tris pH 8.0) and NaCl/Tris solution (150 mM NaCl, 50 mM Tris pH 8.0) in a ratio of 1:3 respectively, for a final concentration of 2 M urea. One μg of mass spectrometry grade trypsin (Trypsin Gold, Promega; 1 mg/mL in 0.03% acetic acid), was added and proteins were trypsinized at 37 °C overnight on a nutator. The next day, the digested peptides were acidified to a final concentration of 0.25% formic acid and 0.25% trifluoroacetic acid. Samples were desalted using a 50 mg C18 Sep-Pak column (Waters), and dried using a SpeedVac vacuum concentrator. Next, samples were resuspended in 90 μL of 65% acetonitrile and 1% formic acid and subjected to hydrophilic interaction chromatography (HILIC) using a TSKgel amide 80 column with a 5 μm particle size (Tosoh Bioscience). Ten fractions were collected at 1-min intervals with the fraction collection gradient starting at 92% solvent A (80% acetonitrile, 0.005% trifluoroacetic acid) and 8% solvent B (0.025% trifluoroacetic acid). The fraction collection gradient ended at 80% solvent A and 20% solvent B. Fractions were dried in a vacuum concentrator and resuspended in a solution of 0.1% trifluoroacetic acid and 0.1 picomole/μL angiotensin II peptide (Sigma-Aldrich).

Each fraction was analyzed using a ThermoFisher Q Exactive HF Orbitrap mass spectrometer coupled to an UltiMate 3000 HPLC system and a C18 capillary column (20 cm length, 125 μm inner diameter) packed in-house with 3-μm, 200-A C18AQ particles (Prontosil). Peptides were separated over a 60-min linear gradient from 8% to 40% acetonitrile in 0.1% formic acid at a flow rate of 200 nL/min. The mass spectrometer was operated in data-dependent mode, and survey scans were acquired in the Orbitrap mass analyzer over the range of 380 to 2,000 m/z with an MS1 mass resolution of 60,000. The maximum ion injection time for the survey scan was 80 ms with a 3e6 automatic gain-control target. Tandem mass spectrometry was performed by selecting up to the 10^th^ most abundant ion (top10 method) with a charge state of 2, 3, or 4 and with an isolation window of 2.0 m/z. Selected ions were fragmented by higher energy collisional dissociation (HCD) with a normalized collision energy of 27, and the tandem mass spectra were acquired in the Orbitrap mass analyzer with a mass resolution of 15,000. For the database search, raw spectra were searched in a SORCERER system (Sage-N Research, Inc.) using SEQUEST software (Thermo Fisher Scientific), and all entries were from the human UniProt proteome database. The following parameters were used in the database search: semitryptic requirement, a mass accuracy of 15 ppm for the precursor ions, a differential modification of 8.0142 Da for lysine and 10.00827 Da for arginine, and a static mass modification of 57.021465 Da for alkylated cysteine residues. The XPRESS software, part of the Trans-Proteomic Pipeline (Seattle Proteome Center), was used to quantify all the identified peptides. A Wilcoxon rank-sum test was used to measure statistical significance.

### Lentivirus production

Lentivirus was produced in HEK 293TN cells, which were seeded two days before transfection to achieve about 60% confluency on the day of transfection with viral plasmids. Packaging plasmids VSVg and PAX2, along with the lentiviral plasmid encoding proteins of interest, were co-transfected into HEK 293TN cells in the ratio of 1:3.1:4.2 using Lipofectamine 2000 overnight in Transfectagro. Fresh growth media was changed the next morning (14–16 h post transfection) to replace Transfectagro. The virus-containing medium was collected 4 times every 12 h starting from 24–30 h post transfection. Collected virus medium was filtered through a 0.45 μm syringe filter and stored at 4 °C for up to 48 h if not used immediately.

### Lentivirus transduction

Most cell lines stably expressing proteins of interest were generated via traditional lentivirus transduction. Cells were seeded on 6-well plates one day before transduction to achieve around 80% confluent on the day of transduction. One well of “kill control” cells was included in the selection process where cells in this well would not be treated with virus. During viral transduction, cells in each well were treated with 1 mL of fresh growth medium, 2 mL of virus-containing medium and 8 μg/mL of polybrene (Sigma-Aldrich). The transduction process was repeated three times every 12 h, and after the last transduction, fresh medium was supplemented for 12 h before the selection process. Cells in 6-well plates were trypsinized, seeded into 10-cm dishes and incubated with appropriated selection drugs until all the kill control cells died. During the selection, drug-containing media was changed every two days. Stable expression of proteins of interest was examined by confocal microscopy or Western blot analysis. Transient virus transduction for exogenous protein expression was achieved by spinfection where indicated. On the day of spinfection, 1 million cells were seeded into each well in 6-well plates and spread evenly, and each well contained 0.5 mL of fresh growth medium, 3.5 mL of virus medium and 8 μg/mL of polybrene. After seeding the cells, the 6-well plates were immediately centrifuged at 930 *g* for 2 h at 33 °C. After centrifugation, without aspirating the transduction medium, 2 mL of fresh growth medium was added to each well, and cells were incubated in a 5% CO_2_ atmosphere at 37 °C until use.

### Cell synchronization

#### S-phase synchronization by double thymidine block

When cells reached 60–70% confluency, 2 mM thymidine was added to the medium for 18 h. Thymidine-containing medium was then discarded, and cells were washed with 1X DPBS (Corning) once and supplemented with fresh growth medium for 9 h. Again, 2 mM thymidine was added and left in medium for 15 h. Cells were washed with 1X DPBS once and released into fresh medium for indicated time before harvest for assays.

#### Prometaphase synchronization by S-trityl-L-cysteine

To synchronize cells in prometaphase, 5 μg/mL of S-trityl-L-cysteine (STLC) was added to confluent dishes of cells for 16 h. Mitotic cells were dislodged from dishes by gentle tapping, washed with 1X DPBS, and collected. Cells pellets were stored at −80 °C if not used immediately.

### Cell cycle analysis

#### Propidium iodide staining and flow cytometry

Measurement of DNA content in fixed cells by propidium iodide staining and flow cytometry was performed as previously described (Rodgers, 2006) with slight modifications. Briefly, experiments were performed in 12-well plates, and cells from each well were lifted by trypsinization and pelleted at 1000 *g* for 3 min. Cell pellets were washed with 1X DPBS once, pelleted again, and resuspended in 100 μL of 1X DPBS. One mL of ethanol prechilled to −20 °C was added to each sample slowly with vortexing, and cells were fixed at 4 °C overnight. Next morning, fixed cells were pelleted at 1000 *g* for 5 min, resuspended in FACS buffer (1% FBS in 1X DPBS), transferred to a 96-well V bottom plate (Greiner Bio-One), and pelleted in the plate. Cells were washed once with FACS buffer and incubated with labeling solution (50 μg/mL propidium iodide and 1 mg/mL RNase A in FACS buffer) for 30 min at 37 °C. The plate was wrapped in foil and stored at 4 °C until samples were analyzed by flow cytometry (BD Accuri C6).

#### Western blot

Cells were lysed with RIPA buffer (150 mM NaCl, 25 mM Tris pH 8.0, 1 mM EDTA, 1% Triton X-100, 0.5% sodium deoxycholate, and 0.1% SDS) supplemented with cOmplete protease inhibitor cocktail (Roche) and phosphatase inhibitors mixture (17.5 mM β-glycerophosphate, 20 mM sodium fluoride, 5 mM sodium pyrophosphate, and 1 mM sodium orthovanadate) on ice. Cell lysates were sonicated for 2 pulses at 20% intensity and centrifuged at 13,000 *g* for 5 min at 4 °C. Protein concentration of the clarified cell lysates was quantified using the BCA assay (Thermo Fisher). Cell lysates were normalized to 2–3 mg/mL, and denatured with 6X Laemmli Buffer (13.3% SDS, 0.067% bromophenol blue, 52.2% glycerol, 67 mM Tris pH 6.8 and 11.1% β-mercaptoethanol) at 95 °C for 5 min. Samples were analyzed by SDS-PAGE and Western blot with detection by chemiluminescence using Clarity Western ECL substrate (Bio-Rad).

#### Live-cell imaging of H2B

Cells were transfected with siRNA duplexes as described above and seeded on 35 mm glass bottom culture dishes (14 mm diameter, #1.5 thickness, Matsunami Glass). Thirty h after siRNA transfection, regular DMEM growth medium was replaced with DMEM without phenol red (Gibco) supplemented with 10% FBS and 1% P/S. Cells were imaged at 37 °C in a 5% CO_2_ environment. H2B-mCherry was excited by the 561 nm laser, and images were acquired every 4 min for 10–12 h to monitor mitosis progression.

### Isolation of endogenous APC/C for in vitro ubiquitination assays

Endogenous APC/C was isolated from HeLa cells synchronized to prometaphase ny STLC as previously described (Oh et al., 2020). Briefly, collected prometaphase HeLa cells were lysed in ubiquitination assay lysis buffer (150 mM NaCl, 5 mM KCl, 1.5 mM MgCl_2_, 20 mM HEPES pH 7.4, 0.1% NP-40) supplemented with cOmplete protease inhibitor cocktail (Roche) and 1 μL of benzonase (Millipore) per 15-cm plate on ice. Cell lysates were vortexed vigorously for 10 s, sonicated with four pulses at 20% intensity and spun at 13,000 *g* for 5 min at 4 °C. Clarified cell lysates were pre-cleared with washed Protein G Sepharose (BioVision) for 1 h with rotation at 4 °C. Protein concentration of the pre-cleared cell lysates was quantified using the BCA assay (Thermo Fisher), and a small fraction of the pre-cleared cell lysates was reserved and normalized as input. The remaining cell lysates were incubated with anti-ANAPC3 antibody (Santa Cruz Biotechnology, 2 μg antibody per mg of input) for 1 h at 4 °C with rotation, and then washed Protein G Sepharose was added to the sample for incubation at 4 °C with rotation for 2 h. The APC/C coupled beads were washed 5 times with ubiquitination assay lysis buffer (at least 10 times of bead volume) before use in the in vitro ubiquitination assay, which was performed immediately after isolation of APC/C on beads.

### In vitro ubiquitination assays

In vitro ubiquitination assays were performed as previously described with slight modifications (Oh et al., 2020). The assays were carried out in a 20 μL reaction volume, and components of complete reactions included: 0.5 μL of 5 μM UBE1 (125 nM final, R&D Systems), 1 μL of 10 μM UBE2S (0.5 μM final, R&D Systems), 1 μL of 10 μM UBE2C (0.5 μM final, R&D Systems), 1 μL of 10 mg/mL His-TEV-ubiquitin (0.5 μg/μL final, prepared as described below), 2 μL of 0.5 mg/mL securin (0.5 μg/μL final, Abcam), 1 μL of 100 μM DTT, 1.5 μL of energy mix (150 mM creatine phosphate, 20 mM ATP, 20 mM MgCl_2_, 2 mM EGTA, pH to 7.5), 2 μL of 10X assay buffer (500 mM NaCl, 100 mM MgCl_2_ and 250 mM Tris pH 7.5), 5 μL of PAC/C coupled resin and 5 μL of 1X PBS. For control reactions that lacked a certain component, the mixture was supplemented accordingly with 1X PBS. Reactions were performed at 30 °C with shaking for 30 min and stopped by addition of 6X Laemmli Buffer and incubation at 95 °C for 5 min. Samples were analyzed by SDS-PAGE and Western blot.

### Protein expression and purification from *E. coli*

To generate 6xHis-TEV-ubiquitin, E. coli BL21-pRosetta2 transformed with pET-His-TEV-Ub plasmid (a gift from Yuxin Mao, Cornell University) was cultured as previously described (Shami Shah et al., 2019). Briefly, a single colony was picked and grown in terrific broth (TB) supplemented with potassium phosphate buffer (0.17 M monobasic potassium phosphate, 0.17 M dibasic potassium phosphate), ampicillin, and chloramphenicol for 14–18 h as a starter culture. 1 L of TB containing phosphate buffer and antibiotics was inoculated with 25 mL of the starter culture and grown for 6–8 h at 37 °C until OD600 was between 2 and 3. The temperature was then decreased to 18 °C for 1 h, and protein expression was induced by adding 0.25 mM isopropylthio-β-galactosidase (IPTG) to the cultures and for at least 18–20 h at 18 °C. Cells were harvested (2100 *g*, 15 min, 4 °C) and stored at −80 °C until use.

Frozen cell pellets were thawed in bacterial lysis buffer (50 mM Tris pH 8.0, 150 mM NaCl, and 0.1 mM phenylmethylsulfonyl fluoride), sonicated, and centrifuged at 31,000 *g* for 45 min at 4 °C. The supernatant was incubated with washed TALON metal affinity resin (Takara) for 1.5 h under nutation at 4 °C. After incubation, the resin with bound protein was washed extensively with 150 mM NaCl, 50 mM Tris pH 8.0 to get rid of non-specifically bound proteins. His-TEV-Ub was eluted by incubation of the washed resin with elution buffer (300 mM imidazole, 150 mM NaCl, 50 mM Tris pH 8.0) for 10 min at 4 °C. Elution was continued until a Bradford assay showed no more protein was eluting from the resin. All the eluant fractions were pooled and buffer-exchanged to 150 mM NaCl, 50 mM Tris pH 8.0 to remove most of the imidazole in the eluant, concentrated in 3K Amicon concentrators (Millipore), quantified using a Bradford assay, and flash frozen for storage at −80°C until use. Samples were analyzed by SDS-PAGE to verify yield and purity. In some instances, His-TEV-Ub was further purified by size exclusion chromatography with 150 mM NaCl, 20 mM Tris pH 7.5, with fractions collected based on molecular weight and verified by SDS-PAGE, followed by concentration and flash freezing for storage at −80°C.

### Proximity biotinylation by MT-TurboIDs

HeLa cells transfected with pCDNA3-DCX-TurboID-V5 or pCDNA3-EMTB-TurboID-V5 for 30 h were incubated with 500 μM biotin for 10 min at 37 °C under 5% CO_2_. For localization by immunofluorescence, cells were rinsed five times with 1X PBS and processed for immunofluorescence analysis as described earlier (see Confocal microscopy section). For streptavidin-agarose pulldown and Western blot, cells were rinsed five times with 1X PBS and lysed with RIPA buffer supplemented with cOmplete protease inhibitor cocktail (Roche). Cell lysates were sonicated with four pulses at 20% intensity and centrifuged at 13,000 *g* for 5 min at 4 °C. Protein concentration of the clarified cell lysates was quantified using the BCA assay (Thermo Fisher), and a small fraction of the clarified cell lysates was reserved and normalized as input. The remaining cell lysates were subjected to immunoprecipitation using streptavidin-agarose (Thermo Fisher) with rotation at 4 °C overnight. The resin was then centrifuged for 5 min at 1000 *g*, washed two times with RIPA buffer, one time with 1 M KCl, one time with 0.1 M Na2CO3, one time with 2 M urea in 10 mM Tris pH 8.0, and two times with RIPA buffer to eliminate non-specific binding. Samples were then denatured and analyzed by SDS-PAGE and Western blot with detection by chemiluminescence using Clarity Western ECL substrate.

### Statistical analysis

For all experiments involving quantification, statistical significance was calculated using an unpaired two-tailed Student’s t-test with unequal variance in Microsoft Excel, a Mann-Whitney U test in GraphPad Prism, or a one-way ANOVA with Tukey post-hoc test in GraphPad Prism as indicated in the figure legend. Statistical significance of p < 0.05 or lower is reported, and the number of biological replicates analyzed is stated in the legend. All the raw data were plotted into graphs using GraphPad Prism. In figures containing scatter plots with straight connecting lines between markers to show changes in measurement over time, the mean at each timepoint was plotted, and the error bars represent standard deviation. For regular scatter plots, the black line indicates the mean, and each dot represents an individual biological replicate.

## ACKNOWLEDGMENTS

We acknowledge support from the NIH (J.M.B.: R01GM131101; M.B.S.: R35GM141159) and the Sloan Foundation (J.M.B.: Sloan Research Fellowship). We thank Reika Tei, Wendy Cao, and Riasat Zaman for technical assistance, the Bretscher lab for the HEK 293TN cell line, the Yu lab for the PLEKHA5 cDNA, the Lammerding lab for the pCDH-CMV-MCS-EF1α-Puro plasmid, the Fromme and Emr labs for equipment, and Michael Goldberg and members of the Baskin lab for helpful discussions.

## AUTHOR CONTRIBUTIONS

Conceptualization: X.C., A.S., J.M.B.; Funding Acquisition: M.B.S., J.M.B.; Investigation: X.C., A.S., E.J.S.; Project Administration: M.B.S., J.M.B.; Supervision: M.B.S., J.M.B.; Writing – original draft: X.C., J.M.B.; Writing – review & editing: X.C., A.S., E.J.S., M.B.S., J.M.B.

