## Supplemental Information for "Proximity labeling reveals spatial regulation of the anaphase-promoting complex/cyclosome by a microtubule adaptor"

**Contents:**

Figures S1–S5
Legend for Table S1

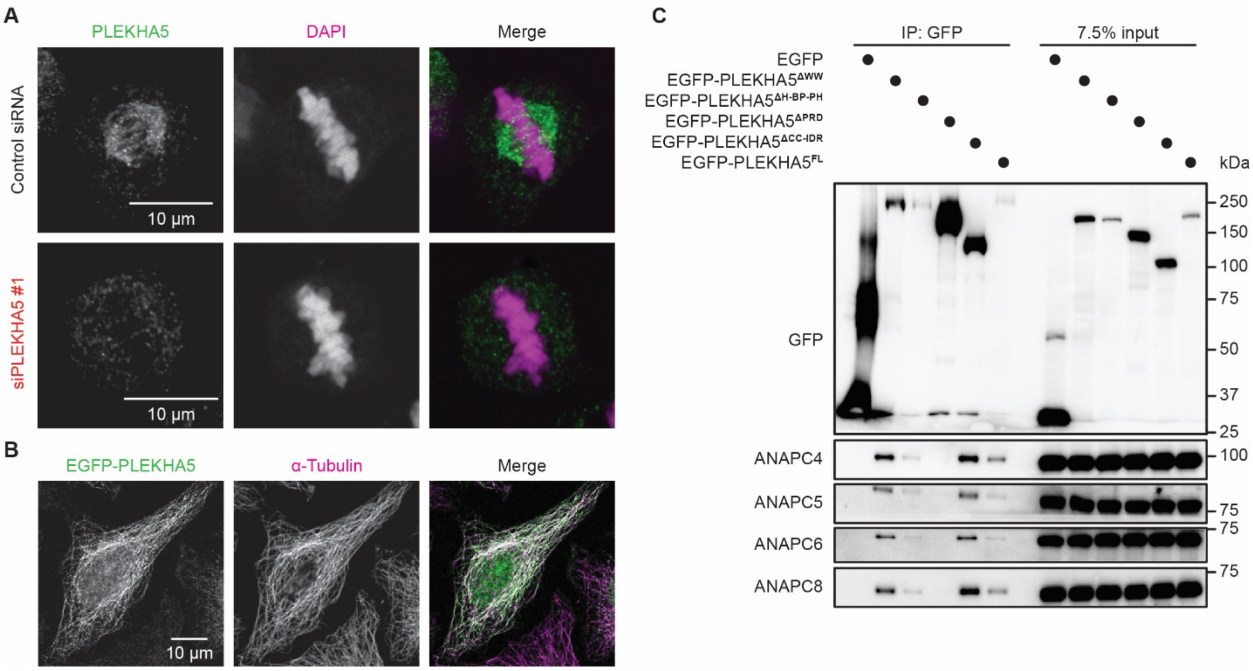


**Figure S1 (related to Figure 1). Localization of PLEKHA5 to microtubule cytoskeleton and domain mapping of PLEKHA5–APC/C interaction.** (A) Immunofluorescence of endogenous PLEKHA5 in HeLa cells transfected with an siRNA against PLEKHA5 (siPLEKHA5 #1) or a control siRNA. Signal of PLEKHA5 (green) diminishes upon siPLEKHA5 #1 treatment, demonstrating the efficiency of knockdown and specificity of the antibody against PLEKHA5. (B) Colocalization of GFP-PLEKHA5 with microtubules as detected by anti-α-tubulin immunofluorescence. (C) The proline-rich domain (PRD) of PLEKHA5 is necessary for the interaction with APC/C. Shown is western blot analysis of α-GFP IP from HeLa cells transfected with GFP or the indicated GFP-PLEKHA5 truncation constructs and blotted for endogenous APC/C subunits.

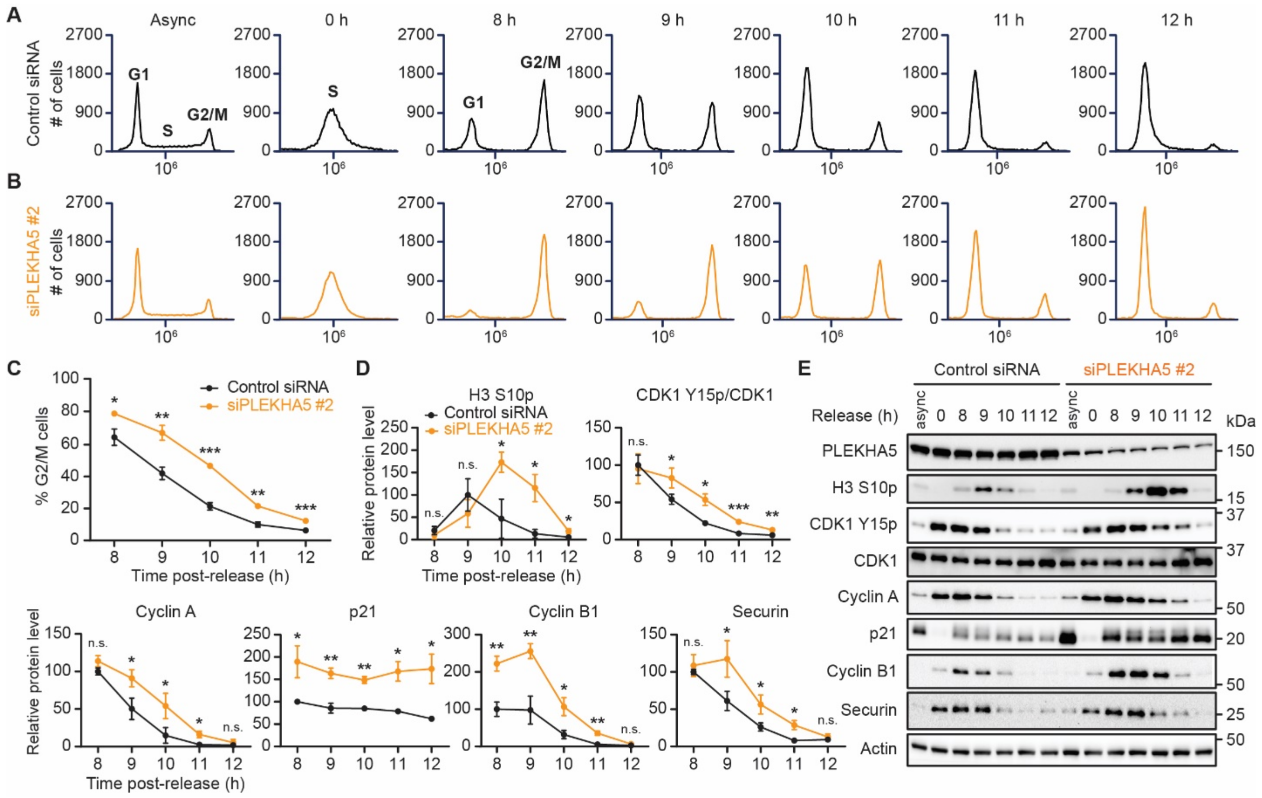


**Figure S2 (related to Figure 2). PLEKHA5 knockdown with siRNA #2 causes accumulation of cells in G2/M assessed by flow cytometry and Western blot.** (A-C) siPLEKHA5 #2 treatment leads to accumulation of HeLa cells in G2/M phase. DNA content of asynchronous (async) or DTB synchronized HeLa cells treated with the control siRNA (A) or siPLEKHA5 #2 (B) were evaluated by propidium iodide staining and flow cytometry analysis, and the percentage of cells in G2/M phase were quantified and shown in (C) (n=3). (D-E) G2/M protein markers and APC/C substrates persists in cells treated with siPLEKHA5 #2. Shown are representative western blots (E) and quantifications of protein levels (D) in lysates from DTB synchronized HeLa cells subject to control siRNA or siPLEKHA5 #2 (n=3). Student's t-test: n.s. not significant; * $p < 0.05$; ** $p < 0.01$; *** $p<0.001$.

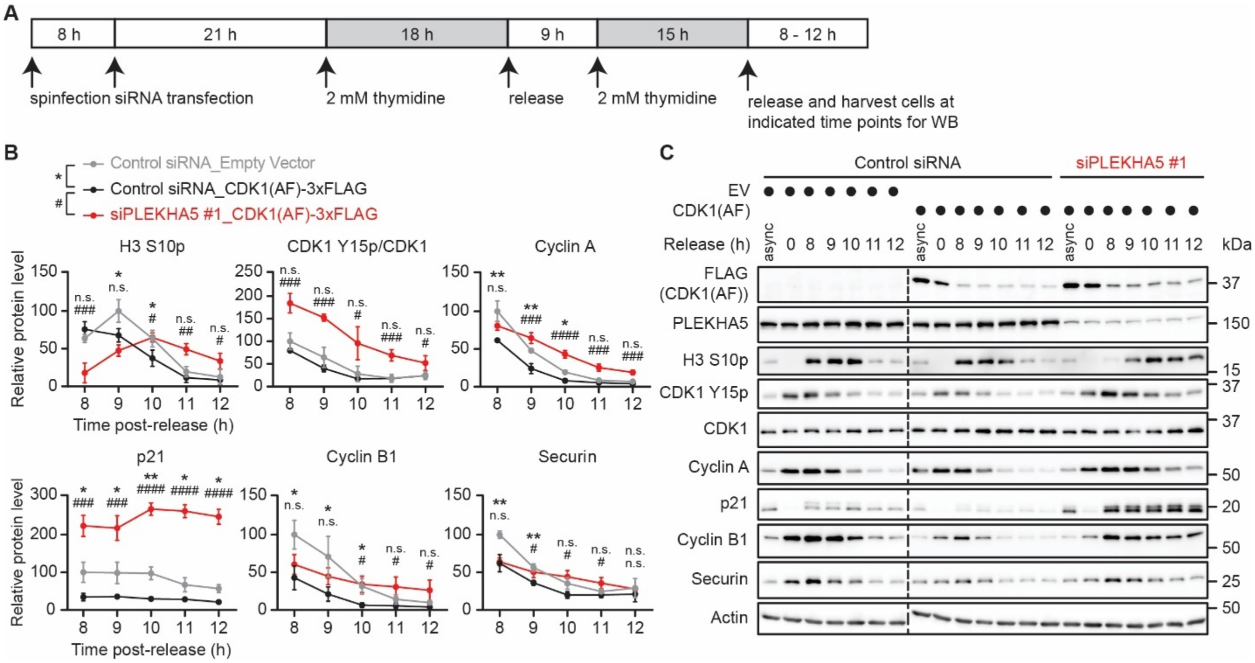


**Figure S3 (related to Figure 3). Knockdown of PLEKHA5 results in buildup of mitosis markers and APC/C substrates in S phase-synchronized HeLa cells expressing a phosphodeficient CDK1 T14A/Y15F (AF) mutant that allows partial G2/M checkpoint bypass.** (A) Schematic representation of experimental timeline. HeLa cells were transduced with lentivirus encoding CDK1(AF)-3xFLAG or empty virus (EV) via spinfection. Eight h after the spinfection, cells were treated with an siRNA targeting PLEKHA5 or a control siRNA and subjected to DTB. Cells were rinsed and released into fresh medium for indicated amount of time before harvest for Western blot analysis. (B) Plots show quantification of the levels of protein markers analyzed by Western blot in (C) (n=3). ANOVA (one-way, Tukey): n.s. not significant; * and # $p < 0.05$; ** and ## $p < 0.01$; *** and ### $p<0.001$; **** and #### $p < 0.0001$.

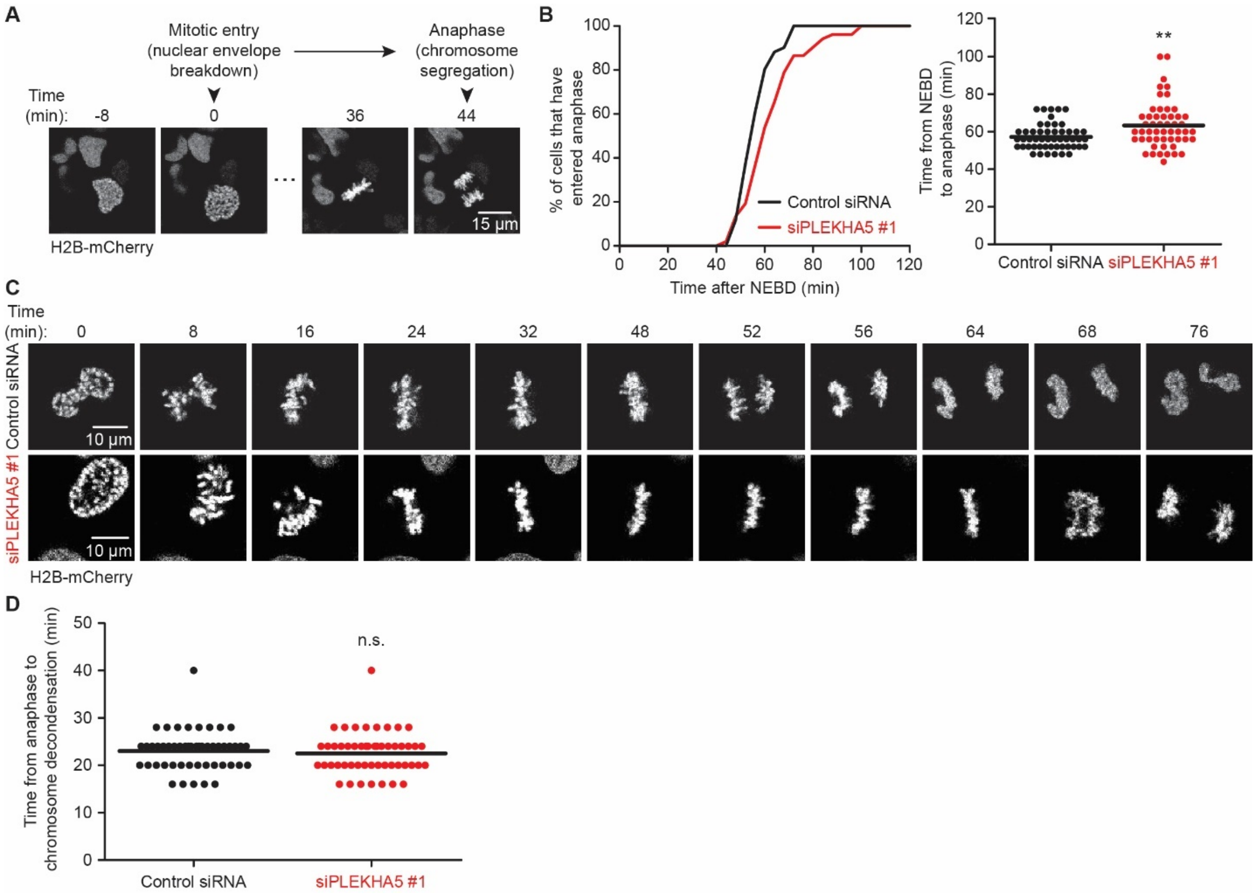


**Figure S4 (related to Figure 3). Mitotic progression analysis in asynchronous HeLa cells by time-lapse, live-cell imaging of H2B-mCherry.** Asynchronous HeLa cells stably expressing H2B-mCherry were transfected with an siRNA targeting PLEKHA5 or a control siRNA. Thirty h after transfection, cells were imaged at 4-min intervals for 10–12 h. (A) Schematic representation of experimental design and quantification. The time from nuclear envelope breakdown (NEBD) to anaphase onset (chromosomal segregation) allows for quantitation of mitosis progression in single cells. (B) Quantification of mitosis progression in asynchronous HeLa cells, with time from NEBD to anaphase plotted as cumulative frequency (left) and scatter plot (right) (n=50–51 cells). (C) Representative series of still images from a time-lapse movie of the nucleus in asynchronous, H2B-mCherry-expressing HeLa cells treated with a control siRNA for siPLEKHA5 #1 used for quantification in (B). Numbers on top of the images are the time from nuclear envelope breakdown (t = 0 min) until the images were taken. (D) PLEKHA5 knockdown did not affect mitotic exit. Quantification of mitotic exit in asynchronous HeLa cells, with time from anaphase to chromosome decondensation plotted as scatter plot (n=50–51 cells). Plotted results were from two separate experiments. Mann-Whitney U test: n.s. not significant; ** $p < 0.01$.

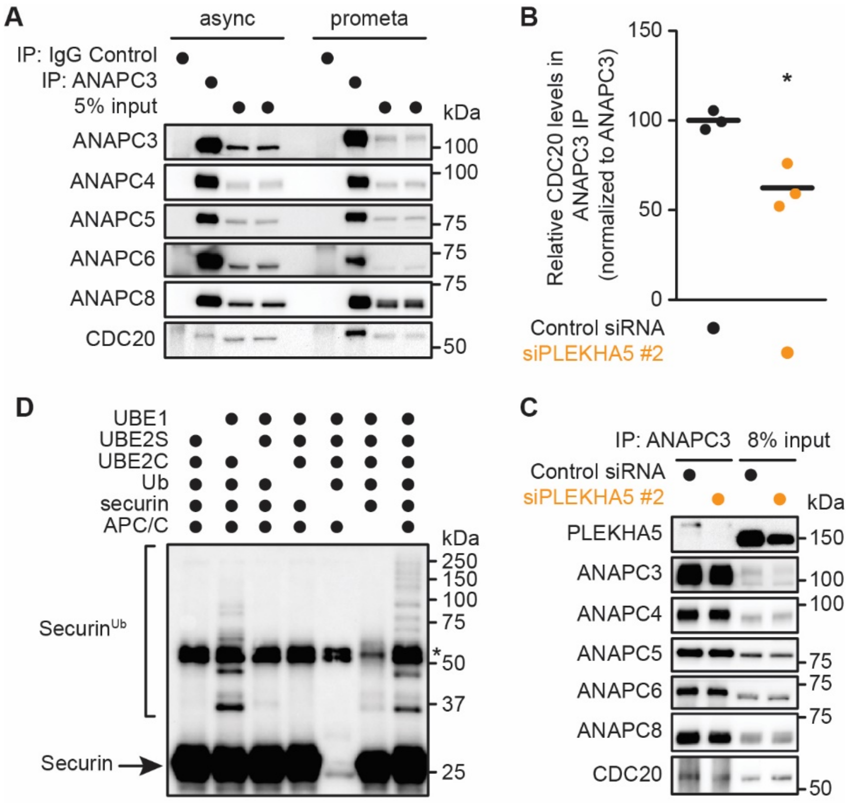


**Figure S5 (related to Figure 4). Effect of PLEKHA5 knockdown on APC/C composition and in vitro E3 ligase activity.** (A) Endogenous APC/C can be purified using α-ANAPC3 IP from cells that were either asynchronous (async) or synchronized to prometaphase (prometa) by treatment with 5 µg/mL STLC for 16 h. Show is western blot characterization of APC/C enriched from HeLa cells compared to whole cell lysate (input). (B-C) siPLEKHA5 #2 causes a decrease in APC/C association with its key mitotic CDC20. Shown are representative western blot analysis of co-IP experiments for endogenous ANAPC3 (C) and quantification of CDC20 levels in the IPs (B) (n=3). (D) APC/C was purified from HeLa cells by incubation of lysates (pre-cleared with protein G Sepharose) with α-ANAPC3 antibody and then protein G Sepharose, followed by rinses to yield APC/C on resin. In vitro ubiquitination assays with the APC/C substrate securin were set up by addition of E1 (UBE1), two E2s (UBE2S and UBE2C), ubiquitin (Ub), ATP, and APC/C (on resin), at 30 ºC for 30 min with gentle agitation, followed by denaturation and Western blot, probing for securin. Indicated on the blot is non-ubiquitinated securin (~25 kDa), mono- and poly-ubiquitinated securin (securin$^{Ub}$), and, with an asterisk (~55 kDa), IgG heavy chain from the APC/C purification. Note appearance of securin$^{Ub}$ only lane 7, which contains all reaction components (as well as some in lane 2, containing UBE2C, which supports multi-monoubiquitination and/or generation of short ubiquitin chains, even in the absence of UBE2S, which generates longer polyubiquitin chains). Student's t-test: * $p < 0.05$.

**Table S1. Raw data from SILAC mass spectrometry experiments** (separate .xlsx file).
